# Synonymous codon usage defines functional gene families

**DOI:** 10.1101/2024.10.31.621346

**Authors:** Farzan Ghanegolmohammadi, Shinsuke Ohnuki, Shane Byrne, Thomas J. Begley, Peter C. Dedon

**Affiliations:** Department of Biological Engineering, Massachusetts Institute of Technology, Cambridge, Massachusetts, 02139, USA; Department of Integrated Biosciences, Graduate School of Frontier Sciences, The University of Tokyo, Kashiwa, Chiba, 277-8562, Japan; The RNA Institute and Department of Biological Sciences, University at Albany, Albany, NY 12222, USA; Antimicrobial Resistance Interdisciplinary Research Group, Singapore-MIT Alliance for Research and Technology, Singapore

**Keywords:** Codon analytics, codon metric, functional networks, isoacceptor frequency, *Saccharomyces cerevisiae*, synonymous codons, translational regulation

## Abstract

**Background:** The degeneracy of the genetic code is increasingly recognized for roles in regulating translation rate, protein folding, and cell response. However, the functional genomics of codon usage patterns remains poorly defined. We previously showed that prokaryotic and eukaryotic cells respond to individual stresses by uniquely reprogramming the tRNA pool and the dozens of tRNA modifications comprising the tRNA epitranscriptome to cause selective translation of mRNAs from codon-biased genes. Here, we systematically defined distinct values of codon bias in the *Saccharomyces cerevisiae* genome by modeling isoacceptor codon distributions using a new statistical toolbox – analysis of synonymous codon signatures (ASCS).

**Results:** Application of ASCS to the *S. cerevisiae* genome revealed linear relationships between patterns of codon bias and gene function using canonical correlation analysis. By mapping codon-biased open reading frames (ORFs) onto a functional network of Gene Ontology (GO) categories, we identified 115 gene families distinguished by unique codon usage signatures. The codon usage patterns were found to strongly predict functional clusters of genes, such as translational machinery, transcription, and metabolic processes.

**Conclusions:** The ASCS-derived model of codon usage patterns in *S. cerevisiae* reveals functional codon bias signatures and captures more biologically meaningful information when compared to other codon analytical approaches.

## Background

The translation of genetic information into functional proteins involves mapping codons to amino acids, with most amino acids (except Met and Trp) encoded by more than one synonymous codon. Since changing synonymous codons do not alter a protein sequence, they were previously considered to be redundant and their mutations silent. However, there is now overwhelming evidence that the choice of synonymous codons acts a critical regulator of translation efficiency,^1–3^ mRNA stability and splicing,^3^ protein folding,^3,4^ cell fitness,^3,5^ and stress response,^3,6^ with so-called “silent mutations” causing a variety of human diseases.^7^

Despite this knowledge, little is known about the rules governing codon usage patterns genome-wide or the existence of signatures of codon bias affecting functional genomics.^8,9^ For example, we have described a molecular pathway regulating gene expression at the level of translation in which stress-induced reprogramming of 40-50 chemical modifications of transfer RNAs – the tRNA epitranscriptome – and the pool of 45-250 expressed tRNAs to selectively translate mRNAs enriched with codons matching the reprogrammed tRNAs.^10–18^ These codon-biased mRNAs appear to be derived from families of stress response genes, yet these families and their functional relationships have remained elusive.

Effects of codons usage patterns can be computationally estimated using a large number of codon-based metrics. These metrics are primarily mathematical models based on each codon or a subset of codons and are designed to describe a biological phenomenon.^19,20^ They are particularly helpful when experimental data is not available; however, each metric is designed to capture a different biological aspect, so the specific usage of each metric must be considered. Among them, the most straightforward are those that consider the frequency of each codon in a gene relative to all other codons (total codon frequency, TCF) or relative to its synonymous partners (isoacceptor codon frequency; ICF) which leads to their widespread usage.^6,10,12–14,16,17^ For example, we employed ICF to statistically predict the role of Trm9 in enhancing translation of mRNAs enriched in Arg-AGA and Arg-AGG codons.^21^ Due to the centrality of codons usage patterns in cell biology, it is important to develop powerful approaches to quantify the extent of codon bias in combination with other biological modalities such as gene function. This helps reveal novel biological aspects that are not captured with the current metrics.

Here we report a new statistical approach of analysis of synonymous codon signatures (ASCS) that maps codon-biased open reading frames (ORFs) onto a functional network of Gene Ontology (GO) categories. Application of ASCS to the *Saccharomyces cerevisiae* genome reveals that codon usage patterns strongly predict gene function, identified 115 functional gene families distinguished by codon usage signatures, and highlights complex codon patterns that have regulatory potential.

## Results

### Biased use of individual synonymous codons does not define gene function

As shown in the workflow depicted in **Figure S1**, the ASCS approach of quantifying usage patterns of synonymous codons begins with a choice of codon counting metric. Dozens of metrics and indices have been developed to quantify frequencies of synonymous codon usage in the coding sequence (CDS) of a mRNA.^20^ Depending upon the application, such as evolutionary genomics or translation efficiency, each metric varies in the extent of considered features, such as the frequency of a particular amino acid in the protein or comparison to a reference set of genes. Given our goal of quantifying gene-specific patterns of synonymous codon usage across a genome, ICF was chosen as the most appropriate codon usage metric given the evidence for its power to predict protein levels in cellular stress responses^6,10,12–14,16,17^. The analysis starts with calculating the average synonymous codon usage data across the genome. This illustrates both the abundance of specific genetically-encoded amino acids and the frequency of specific synonymous codons. For example, cysteine is a relatively rare two-codon amino acid in yeast, with the encoding TGT more abundant – thus designated “optimal” – than its synonymous partner TGC (**Fig. 1A**). These genome codon usage averages are then used to calculate gene-specific ICF values by dividing the total number of a specific synonymous codon by the genome average usage value. As a convenient way to present ICF data, we have traditionally calculated the Z values for the ICF data, which roughly represents the number of standard deviations that the ICF differs from the genomic mean and specifies over- and under-usage as positive and negative values. While we successfully used this ICF approach to predict synonymous codons associated with stress-induced translation up- and down-regulation,^6,10,12–14,16,17^ the approach makes the simplifying assumption of a normal distribution of codon usage patterns, which is an over-simplification.

**Figure 1.**
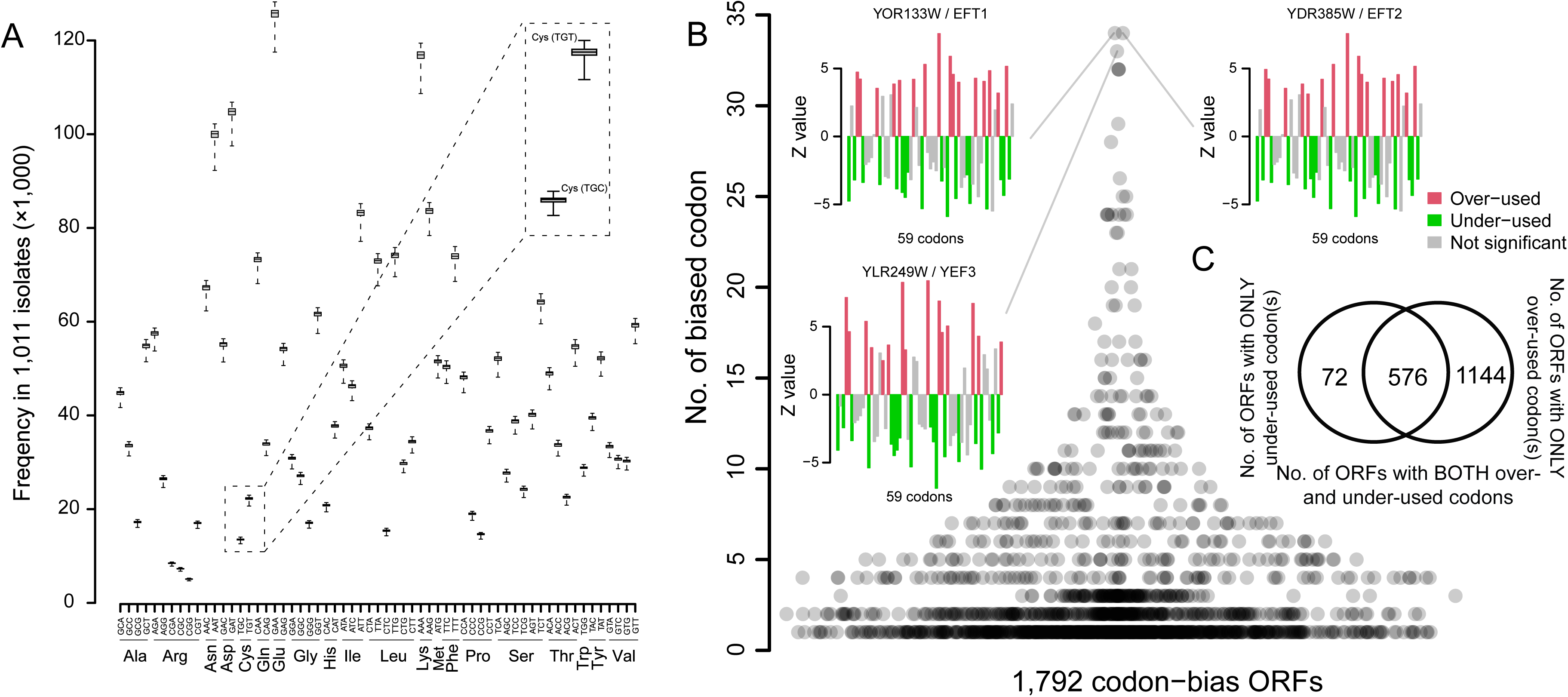
Codon frequency analysis of the *S. cerevisiae* genome. (**A**) Average frequency of synonymous codon usage in 1,011 budding yeast isolates^44^. Data presented as box plots: median, center line; standard deviation, error bars; and upper and lower quartiles, gray boxes. Inset: Codon frequencies of cysteine (Cys) showing preferred isoacceptor (TGT). (**B**) Bee-swarm plot of number of significantly biased codon(s) in codon-biased ORFs (FDR = 0.05). Z values of top 3 codon-biased ORFs are shown. (**C**) Venn diagram of the number of codon-biased ORFs.

As a more rigorous approach, we modeled codon usage data assuming a binomial distribution, converting gene-specific ICF data for 59 codons to Z values using the Wald-test (**Fig. S1**). Following conversion of Z to q values, we then defined codon-biased ORFs as those with at least one codon with a q value significantly different from the genome average (FDR = 0.05). Almost one-third of 5,797 yeast genes were found to be codon biased (1,792 ORFs) as shown in **Table S1**. Of 1,792 codon-biased ORFs, 1,144 and 72 ORFs possess only over- or under-used codons, respectively, while 576 contain both over- and under-used codons (**Fig. 1B,C**). Our previous study had only assessed 425 ORFs that over- and under-used codons based on the assumption of a normal distribution,^21^ with the numerical difference due to our current use of the Wald-test and binomial distribution fit. As expected, highly codon-biased ORFs are mainly involved in protein translation, such as the paralogs *EFT1* and *EFT2* that both catalyze ribosomal translocation during protein synthesis (**Fig. 1B**). Several interesting features of yeast codon usage emerged from this analysis: (1) CGC, CGG, and CGT (Arg), CAC (His), and TCC (Ser) are never under- used in any genes; (2) GGT (Gly) is by far the most frequently used codon (361-fold over-usage); and (3) TTG (Leu) and AAG (Lys) are the next most frequently used codons with 236- and 233-times over-usage, respectively. The 576 ORFs that both over- and under-use specific synonymous codons are mainly responsible for a wide range of functionalities such as metabolic processes and translation (**Table S2**). However, the 1,144 and 72 ORFs possessing either over- or under-used codons, respectively, but not both, were not enriched in any individual GO categories (FDR = 0.05), suggesting that individual codon biases occur across a wide range of functional categories and do not reflect genome-wide function codon signatures. Furthermore, since each of 59 codons was significantly under- or over-used in at least one ORF (**Table S3, Fig. S2A**), we compared the genomic ratio of each codon (**Fig. S2B;** number of a given codon/number of all codons) to its frequency of biased use (how many ORFs showed biased usage of the codon). Again, there were no strong correlations of biased distribution of codons in the genome (**Fig. S2C-E**). For example, the most frequently occurring codons do not necessarily show higher over-usage in ORFs. However, heteroscedasticity (i.e., non-constant standard deviations) of the results (**Fig. S2C-E**) does not allow drawing any general rules. These results reveal that biased use of individual synonymous codons is not a good predictor of gene function, which raises the question of biased use of groups of synonymous codons.

### Simple clustering of biased codons weakly predicts gene function

To test the idea that groups of codons with biased usage predict gene function, we first reduced the dimensionality of the codon bias profile by principal component analysis (PCA; **Fig. S3A**). Gaussian mixture model (GMM) clustering was then applied to the first two PC spaces, which represented a cumulative contribution ratio (CCR) of 38% (**Fig. S3B-C**), codons were grouped into four clusters in which cluster I and IV are mainly A/T-ending codons, and cluster II and III are mainly C/G-ending codons (**Fig. 2A**). These cluster-specific codon-biased ORFs show a clear dichotomy between A/T- and C/G-ending codons in PC2 space (**Fig. 2A**).

**Figure 2.**
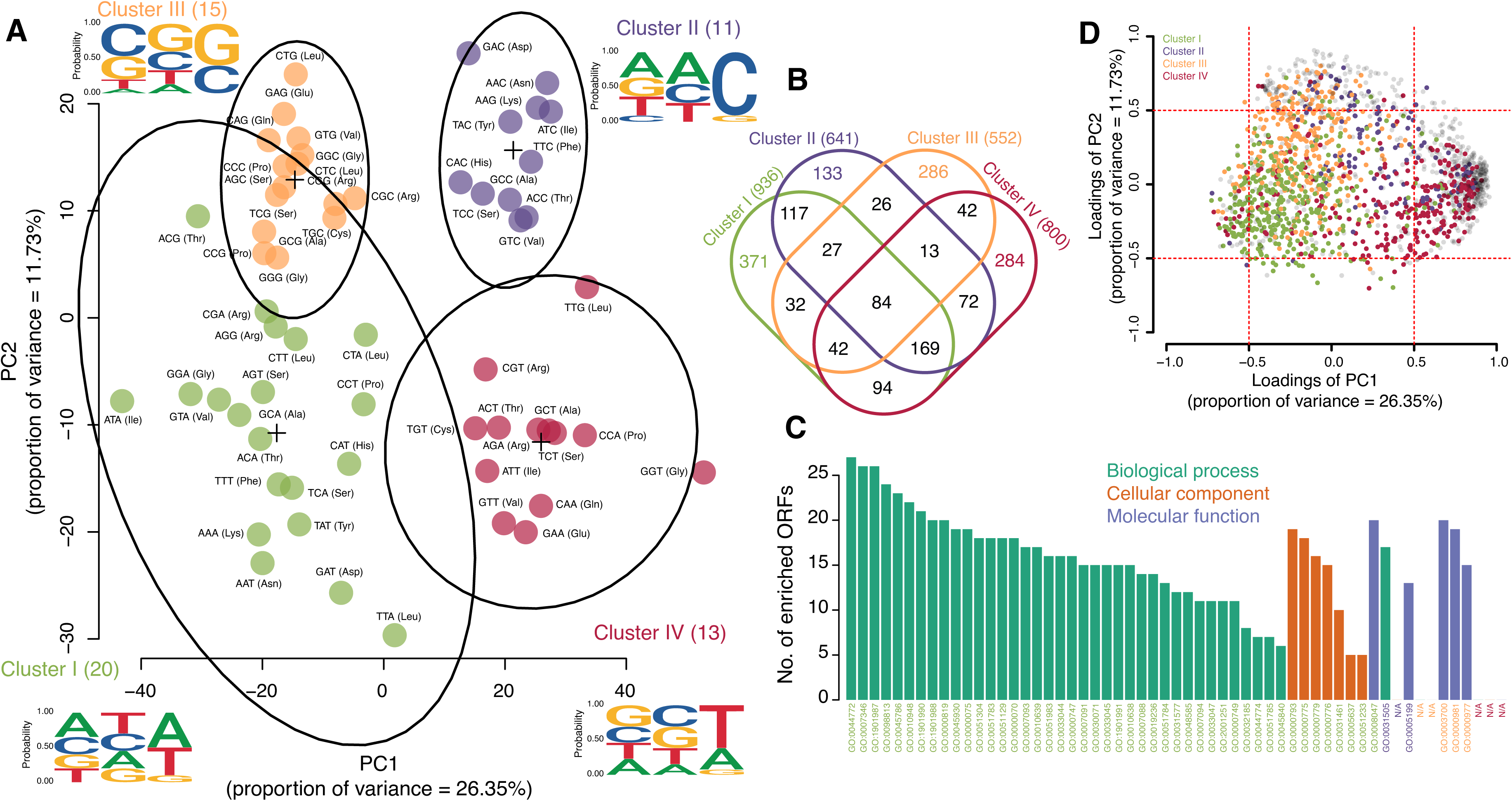
Simple clustering of biased codons weakly predicts gene function. (**A**) A two-dimensional principal component (PC) scores plot illustrates clustering of 59 codons (colored circles). Gaussian mixture modeling of these two PC scores (PC1 and PC2; CCR = 38.09%; **Fig. S3A**) reveled 4 clusters (black circles; I-IV; **Fig. S3B,C**) with unique A/T- or C/G-ending codon patterns (logo views). (**B**) Venn diagram of the number of codon-biased ORFs in each cluster. (**C**) A bar plot of GO enrichment analysis of cluster-specific codon-biased ORFs (color-coded from panel **B**). See **Table S4**. (**D**) Scatter plot of PCA loadings from panel **A**. Each dot represents a codon-biased ORF.

We next used cluster-specific codon-biased ORFs (**Fig. 2B**) for GO enrichment analysis (**Table S4, Fig. 2C**). This revealed that cluster I members are mainly responsible for cell cycle (e.g., GO:0045930) and cell division (e.g., GO:0051783), cluster II are cell wall constituents (GO:0005199) and cell wall organizers (GO:0031505), and cluster III showed transcription regulation activities (e.g., GO:0000981). Members of cluster IV did not enrich to any terms indicating a wide functional variability. However, even though most clusters have specific annotations, most of the members did not play a significant role in forming the clusters (loadings: -0.5<Pearson’s *r*<0.5; **Fig. 2D**). Since this PCA-based approach, which focused on finding linear combinations that account for the most variance in the codon dataset, did not provide a precise or comprehensive overview of the functional map of codon-biased ORFs, we resorted to canonical correlation analysis (CCA) to find linear combinations that account for the most correlation in two datasets.

### Functional gene families are strongly distinguished by signature codon usage patterns

To establish more accurate connections between the codon-biased ORFs and specific gene functions using CCA, we first generated a functional profile of the codon-biased ORFs using GO annotation data and subjected this dataset to hierarchical clustering analysis (**Fig. S4**). Examples of 18 GO functional clusters are shown in **Figure S4** which illustrates the strengths of the relationships among the codon-biased ORFs in each cluster. We then used CCA to obtain linear relationships between the codon bias profiles and the GO profiles. CCA determines a set of orthogonal linear combinations of variables (i.e., canonical variables; CVs) in the codon bias and GO category datasets that best explain the associations between the variables in the two sets. The CCA analysis revealed 11 linear combinations (FDR = 0.05, Bartlett’s chi-squared test; eye- diagram in **Fig. S5**) that maximize the correlation between the codon-bias and functional profiles. These CVs provided spaces for understanding codon signatures in association with gene functions. For example, **Figure 3A** depicts a biplot of the codon-biased canonical variable 1 (cCV1) against the codon-biased canonical variable 2 (cCV2) determined from CCA analysis. This plot shows that cCV1 strongly differentiates regulation of mitotic cell cycle phase transition (cluster 14) and cellular response to nutrient levels (cluster 20) from large (cluster 42) and small (cluster 49) ribosomal subunits; they are well separated along the cCV1 axis (**Fig. 3A,B**).

**Figure 3.**
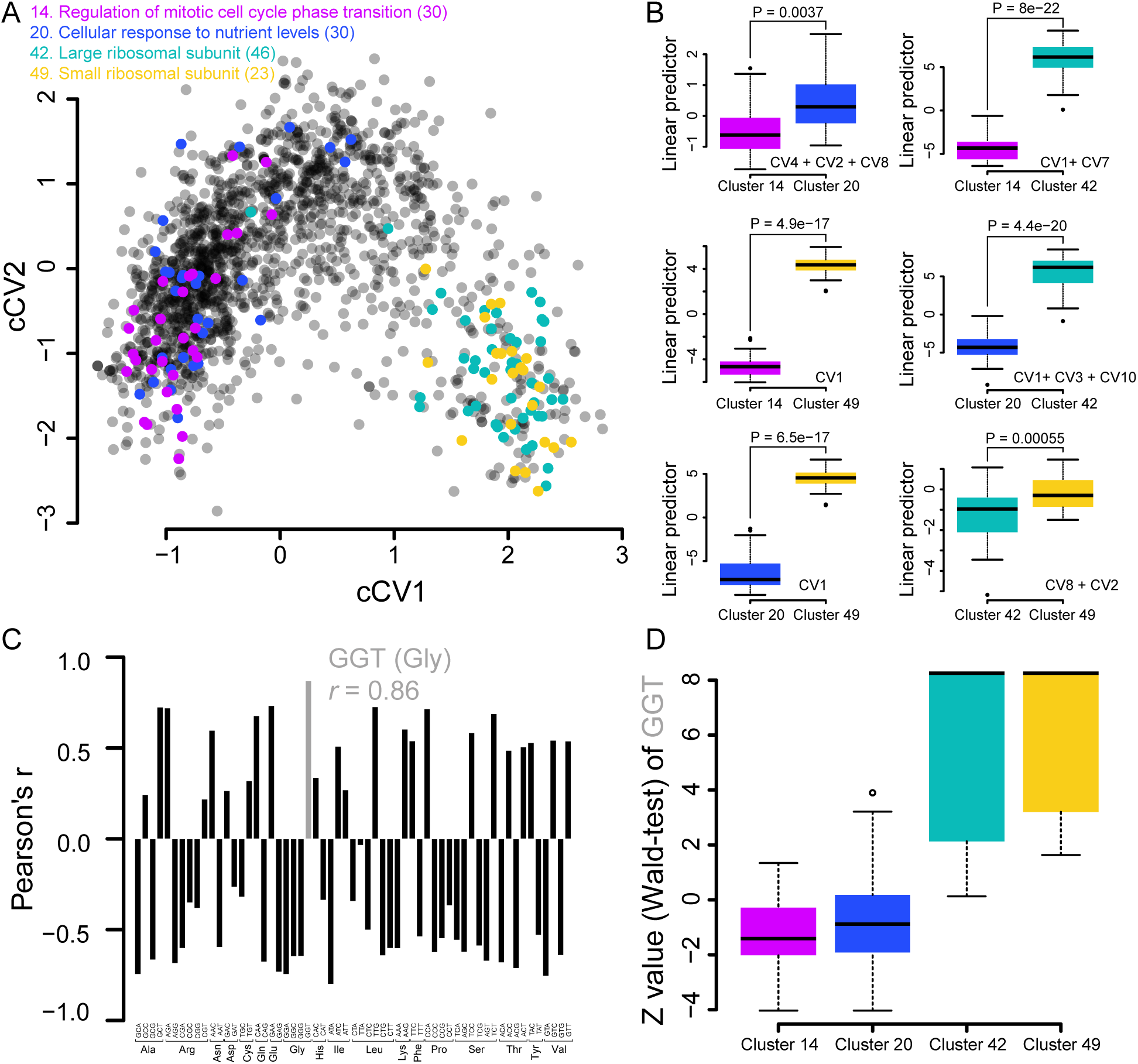
Canonical correlation analysis (CCA) distinguishes functional clusters of codon-biased genes. (**A**) A biplot of the first two codon-biased canonical variables (cCV1, cCV2) as calculated in **Figure S5**. Each colored circle is a codon-biased ORF. Clusters 14, 20, 42, and 49 are color-coded as in **Figure S4**. (**B**) Examples of linear predictors of logistic regression models showing distinctions between functional clusters from panel **A**. Explanatory variable(s) (CVs) of the final model are shown in each case. P values are obtained from a likelihood ratio test of null and optimized models. (**C**) Bar plots of Pearson’s correlation (*r*) analyses of cCV1 and Z values of 59 codons of codon-biased ORFs. The highest *r* value (Gly-GGT) is shown in gray. (**D**) Box and whisker plot of Z values (Wald-test) of GGT (Gly) of the 4 clusters from panel **A**; median, center line; standard deviation, error bars; and upper and lower quartiles, gray boxes.

To systematically define the best combination of CVs that differentiate functional clusters, we employed logistic regression followed by a stepwise approach (see *Methods*). As shown in **Figure S6A**, most clusters are separable by only 2 cCV spaces, indicating clear distinctions caused by codon usage patterns and gene functionalities. The differentiating power of combinations of CVs is illustrated by the fact that cCV1 is not suitable for separating clusters 14 and 20 and clusters 42 and 49, but these clusters are resolved by a combination of cCV2 and cCV8 (**Fig. 3B**). In other examples, 3 CV spaces (cCV1, cCV3, cCV10) are required to separate clusters 20 and 42 (**Figs. 3B, S6B**), while cCV1 and cCV7 are the best separators of clusters 14 and 42 (**Figs. 3B, S6C**).

With functional gene clusters distinguished by the CVs, the next step was to define the patterns of codon usage that account for the observed divisions. Here we estimated the Pearson’s correlation coefficients (*r*; **Fig. S7**) for a comparison of significant CV spaces (11 linear spaces; FDR = 0.05, Bartlett’s chi-squared test; **Fig. S5**) and the estimated Z values of each of 59 codons in the codon-biased ORFs (**Table S6**). As an example, CV1 is strongly correlated with GGT of Gly (*r* = 0.86; **Fig. 3C**). This reflects the codon-usage pattern, estimated as Z values (**Fig. 3D**), in which clusters 14 and 20 mainly under-used GGT but clusters 42 and 49 mainly over-used it. This indicates successful application of CCA in capturing codon-usage patterns and suggests that signatures of biased codon usage are linked to gene functions.

A global network of codon-biased gene families was constructed by estimating Pearson’s correlations between every pair of codon-biased ORFs using CV scores. The resulting correlations were then used to systematically map codon-biased ORFs belonging to 115 functional clusters (**Table S5**). As shown in **Figure 4**, this approach provides a global view of the functional relationships among codon-biased ORFs. In this network, we observed 10 “core” gene clusters (124 ORFs; **Fig. S8A**), which have more significant associations with other clusters (≥ 11) and are directly or indirectly connected to other clusters (**Fig. S9A**), such as mitotic sister chromatid segregation (GO:0000070; 8 members) and mitochondrial translation (GO:0032543; 14 members). These clusters are hubs in the decentralized network. However, hierarchical cluster analysis of significant correlations revealed that these core clusters are not tightly clustered (**Fig. S9B**) implying that all groups, even groups with few associations, play important roles in forming the network’s topology through their interconnections. We also found 63 “dense” gene clusters (1307 ORFs; **Fig. S8B**), defined as ORFs that are <0.1 distance units from the center of the gene cluster. These indicate that majority of the ORFs with similar patterns of codon bias are functionally related genes. Four clusters served as both core and dense clusters (75 ORFs; **Fig. S8C**). From among 73 strongly differentiated codon-biased gene families, **Figure 4** illustrates the granularity of functional distinctions with 17 dense clusters (red dots in **Fig. S8C**) and 1 core+dense cluster (alcohol metabolic processes; turquoise dots in **Fig. S8C**).

**Figure 4.**
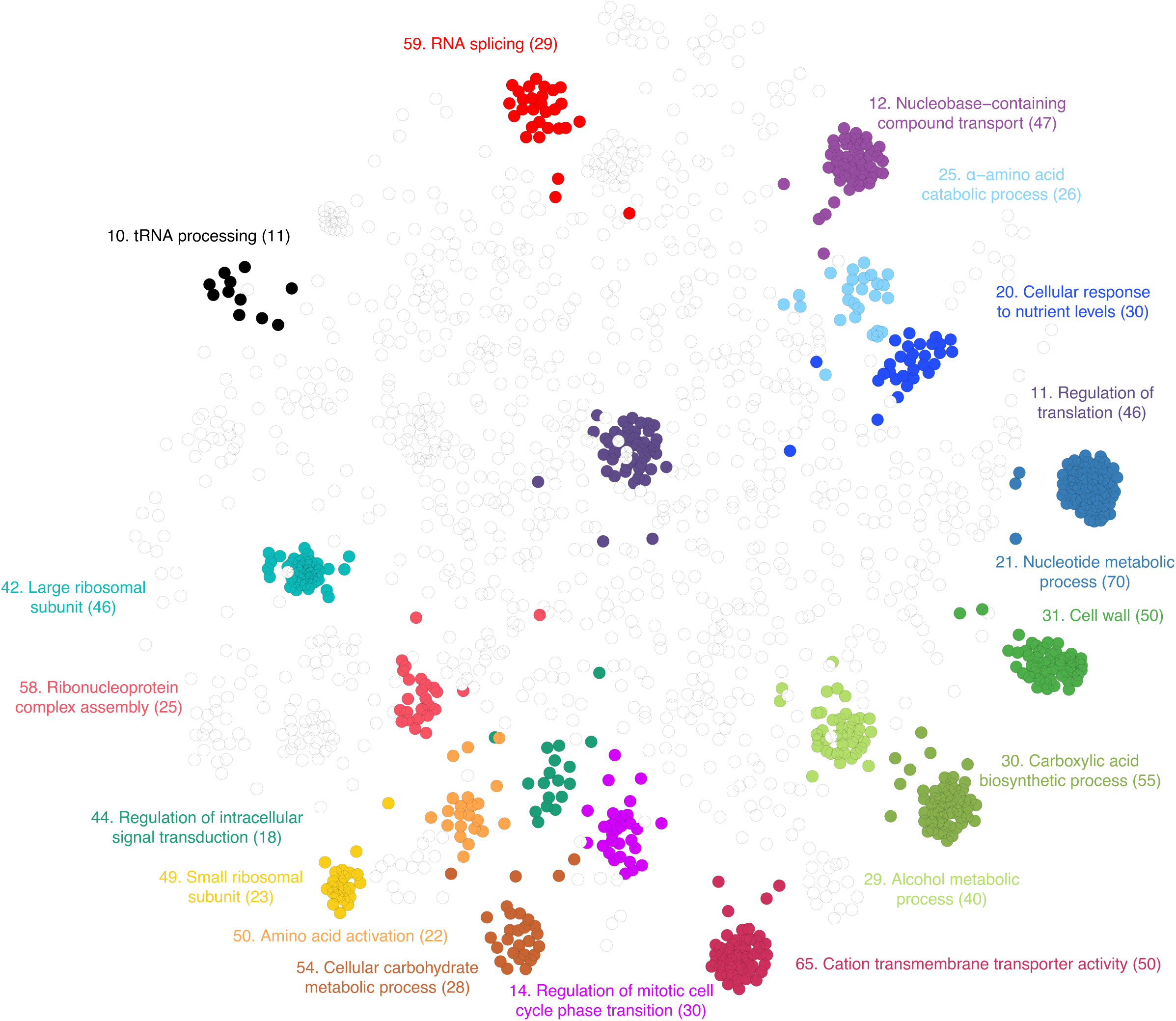
Signature codon usage patterns define functional gene families. A plot (spring layout) of Pearson’s correlations (*r*) between every pair of codon-biased ORFs using the 11 cCV scores reveals strong clustering of codon-biased ORFs and their functions. Of the 115 total clusters detected using this approach, 18 are displayed with corresponding cluster numbers (**Table S5**). Numbers in parentheses represent the number of ORFs in the cluster.

### Evaluating functional information gain

To assess the rigor of our approach, we benchmarked it against our previous work with a simpler codon counting algorithm^21^ and against another codon analytical metric, relative synonymous codon usage (RSCU),^22^ with the former assuming a normal distribution of biased codon usage. Compared to both approaches, our model revealed more codon-biased ORFs (**Fig. S10A**), largely due to the use of more rigorous modeling with a binomial distribution. To assess the contribution of relevant biological information to the performance, as opposed to modeling methodology, we further assessed the degree of similarity between codon-biased ORFs and their functional predictive power for the three approaches, as previously described.^23^ The results show a higher precision and recall curve for GO annotations in this study and support the idea that that our model has higher predictive power for gene function based on a codon bias profile (**Fig. S10B**). In addition to enhancing our understanding of codon bias signatures in functional groups of genes, the observation of 1,792 codon-biased ORFs led to a more comprehensive network revealing functional relationships among codon-biased gene families and a better resolution of gene function (**Fig. S10C, D**).

## Discussion

Here we analyzed codon bias in the *S. cerevisiae* genome using a new statistical toolbox termed ASCS and discovered linear relationships between patterns of codon usage and functional families of genes. These results highlight the fact that, while large biological datasets are typically noisy, a rigorous analytical approach can yield precise estimations limited loss of valuable data and thus capture more relevant biological information. Coupling ICF with statistical modeling approaches that reflect the non-normal distribution of synonymous codons and applying logical strategies to extract relevant biological information, we found that codon-biased genes account for roughly 30% of the yeast genome. Our more inclusive approach reveals that combinations and patterns of codon usage more accurately predict gene function than individual codons. Using this approach, we identified 151 more codon-biased genes (576/5801 = 9.9%) than we previously found using simpler approaches (425/5780 = 7.4%).^21^

Many previous studies have identified codon biases in functional groups of genes, including translation machinery,^9^ protein folding,^24^ and signal transduction.^25^ These functions stand out due to the very high levels of codon bias in the genes involved in these functions. We observed the same biases and gene families, but we found that it is a relatively small group of genes that have these extreme codon bias patterns (**Table S7**) and not surprisingly they are mainly enriched in transcription, translation, and cell wall functions (**Table S5**; effect size: median of sum of absolute Z values of members in each cluster). One could argue that protein synthesis is a vital and energetically expensive process within a cell,^26^ with strong natural selection driving codon usage patterns for efficient translation.

Our ASCS approach, however, provided a more holistic picture of codon-biased gene families. The 115 functional gene families distinguished by codon usage signatures provide a more granular basis for translation regulation of gene expression that could enable cells to regulate molecular and cellular processes under wide range of conditions.^6^ For example, compared to ribosomal subunit (cluster 42 and 49), we found less extreme but still significant codon bias patterns in gene families for alcohol (cluster 29) and sulfur (cluster 8) metabolic processes, with effect size^†1^ of 86.9 and 82.9, respectively. These functional clusters are mainly responsible for cell survival under changing environmental conditions in which selective translation of adaptation proteins is required for a quick and efficient response. Indeed, we and others have observed many examples of selective translation of codon-biased mRNAs from stress-response genes, including the yeast response to oxidative stress,^10,11^ the mycobacterial response to hypoxia and starvation,^12,16,27^ human B-cell production of antibodies,^28^ and the human cell response to toxic exposures.^17^ In all cases, the codon-biased translation is linked to stress-induced reprogramming of the pool of tRNA isoacceptors and the dozens of tRNA modifications comprising the epitranscriptome, with the reprogrammed tRNAs selectively reading specific synonymous codons enriched in mRNAs from families of survival genes. Selective translation of families of codon-biased genes also arises dysfunctionally in many human diseases, such as over-expression of METTL1 and ELP3 in subsets of several cancers causing hypermodification of specific tRNAs and over-translation of codon-biased mRNAs for genes driving cell growth.^29,30^ The revelation of codon bias signatures that define 115 functional gene families generalizes these focused observations, validates a general model of tRNA reprogramming and codon-biased translation,^31^ and opens the door to linking the epitranscriptome to a broader network of environmentally-responsive gene expression.

Our analysis of the *S. cerevisiae* genome using ASCS provides an opportunity to more deeply characterize and validate the role of codon-biased gene families in the yeast translational response to stress. For example, Trm9 installs wobble (position 34) mcm^5^U in yeast tRNA isoacceptors Arg-<u>U</u>CU and Gly-<u>U</u>CC and, with 2-thiolation by CTU1/CTU2, wobble mcm^5^s^2^U in tRNAs Glu-<u>U</u>UC, Gln-<u>U</u>UG, and Lys-<u>U</u>UU.^21,32^ These tRNAs are involved in the codon-biased yeast translational response to exposure to alkylating agents We previously showed that loss of Trm9 led to sensitivity to alkylating agents as a result of reduced translation of mRNAs enriched in Arg-AGA and Arg-AGG codons, including those encoding translation elongation factor 3 (*YEF3*) and the ribonucleotide reductase (*RNR1* and *RNR3*) large subunits involved in the DNA damage response pathway.^21^ Additionally, we identified 422 other genes enriched with combinations of AGA and AGG, as well as enriched with codons considered low-usage codons, including Asp-GAC, Ile-ATC, Tyr-TAC, Lys-AAG, and Phe-TTC.^21^ Importantly, we found that Trm9-dependent codons Arg-AGA and Gly-GAA cluster along the length of genes,^14^ which adds another layer of codon-based regulatory control over translation. The 115 gene families defined with ASCS here provide an opportunity to granulate the yeast phenotypes associated with mcm^5^U, mcm^5^s^2^U, and a variety of other tRNA modifications.

ASCS-defined gene families also explain the yeast response to oxidative stress. We previously reported that UUG-rich mRNAs are selectively translated in yeast cells exposed to hydrogen peroxide (H_2_O_2_)^6^. Along with Lys-AAG, Leu-TTG plays a major role in defining the codon bias signatures of clusters 42 (large ribosomal subunit) and 49 (small ribosomal subunit) (**Fig. 5A**). Most of the 33 up-regulated proteins in the H_2_O_2_ study were revealed as codon-biased in our analysis, 75% over-using TTG and 12 of them mapping to clusters 42 and 49 (**Fig. S11**). Among the up-regulated proteins were *RPL4B*, *RPL2A*, *RPL8A*, and *RPL7B,* which shared the highest codon bias profile similarities in cluster 42 (two-sided t-test, FDR = 0.01; **Fig. 5B**). These genes over-use Lys-AAG, Leu-TTG, Asn-AAC, and Gly-GGT, while they under-use Lys-AAA. Similarly, the highest codon bias profile similarity in cluster 49 were between *RPS13* with *RPS22A*, and *RPS4A* with *RPS23A* (two-sided t-test, FDR = 0.01; **Fig. 5B**), all of which were up- regulated in H_2_O_2_-exposed yeast and over-use Lys-AAG and Leu-TTG. While all of these ribosomal proteins were up-regulated in H_2_O_2_-exposed yeast, their paralogous counterparts (*RPL4A*, *RPL2B*, *RPL8B*, *RPL7A*, *RPS22B*, *RPS4B, RPS23B*) were not detected.^6^ These results, coupled with the observation that proteins related to translation are significantly upregulated by oxidative stress,^6^ further support the idea of selective expression of codon-biased gene families in cell stress responses. The presence of multiple codon enrichments also suggests that this translational regulation of stress response involves biases in more than a single codon and supports the idea that groups of codons are involved. Indeed, >70% of proteins up-regulated by H_2_O_2_ are enriched in both AAG and TTG codons in *S. cerevisiae*.^6^ While TTG has experimentally been proven important in the H_2_O_2_ stress response in budding yeast,^6^ AAG has not been experimentally validated. Interestingly, the other codon for Lys, AAA, is enriched in genes critical for surviving H_2_O_2_ stress in fission yeast, *S. pombe*.^33^

**Figure 5.**
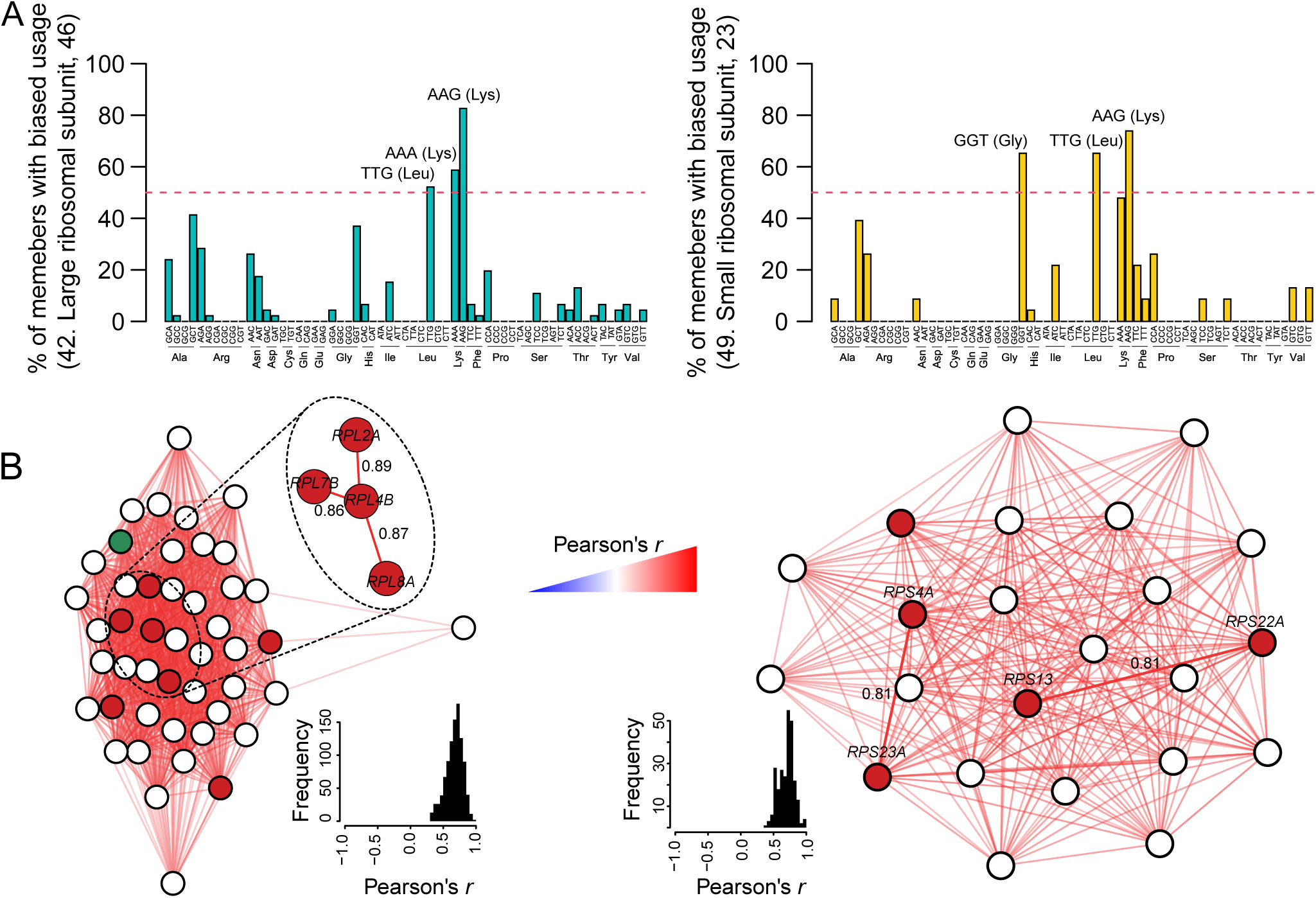
Patterns of biased codon usage in ribosomal genes that have been linked to H_2_O_2_ stress response genes in *S. cerevisiae*. (**A**) Bar plots of percentage of cluster members with biased usage of each codon (over- and under-use) in clusters 42 (large ribosomal subunit; 46 members) and 49 (small ribosomal subunit; 23 members). (**B**) Codon bias profile similarity networks in clusters 42 (left) and 49 (right) using a spring layout. Genes for up- and down-regulated proteins under H_2_O_2_ stress^6^ are shown in red and green nodes, respectively. Edges are *r* values obtained by Pearson’s correlation analysis of the PC scores (CCR = 99%) after applying PCA to Z values. Histograms show distributions of *r* values in each network. Gene with highest profile similarities are shown.

These studies of yeast ribosomal proteins also provided insights into the limits of stress-specific codon-biased translation. Of 79 ribosomal proteins in *S. cerevisiae*, 59 have paralogous partners encoded by different genes. As noted earlier, *RPL22A* possessed a higher enrichment of TTG than *RPL22B*, was selectively upregulated by oxidative stress, and was critical to the *RPL22A* while *RPL22B* was not. Interestingly, a plot of codon bias similarity using Pearson’s correlation (*r*) analysis of the Z values (**Table S1B**) *versus* percentage of sequence identify (SI) using pairwise alignment revealed that higher percentages of sequence identify lead to more similar codon bias profiles (**Fig. S12**). This suggests that there is a “sweet spot” for differing codon biases among the paralogs that will correlate with their differential regulation of cell phenotype.

While there is often a tendency to treat biological data as normally distributed, perhaps due to its simple mathematical and computational specifications, this is not always true.^34^ Indeed, to detect subtle differences in large biological datasets, higher statistical power is required. Here, we used a generalized linear model (GLM) approach, including fitting a binomial distribution to isoacceptor frequencies, to provide clear distinctions among codon usage patterns in genes. The resulting identification of codon-biased gene families provides a higher predictive power for linking cell phenotype to stress-induced codon-biased translation.

## Methods

### Generating the codon bias profile

A summary of the workflow involved in ASCS quantification of codon usage biases in yeast gene families is detailed in **Figure S1**. The process starts with quantifying codon usage in genomic open reading frames (ORFs) in *Saccharomyces cerevisiae* S288C as a reference strain. ORFs were downloaded from The *Saccharomyces* Genome Database (SGD; https://www.yeastgenome.org/) and blocked reading frames and transposable elements were removed (**Table S1A**). The metric for codon usage involved the isoacceptor codon frequency (ICF), which has previously involved quantifying proportions of discrete isoacceptor codon counts in each ORF compared to a genome average using generalized linear models.^35^ However, instead of the original simplifying assumption of a normal distribution of codon frequencies, we used a more rigorous binomial distribution (BI) to fit to the genomic distribution of each of 59 codons (ATG of Met, TGG of Trp, and stop codons [TAA/TAG/TGA] were excluded). The gamlss function of the gamlss package ^36^ was used to accommodate the statistical model as follows:

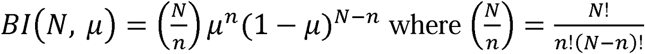

Where, *n* is number of occurrences of a given codon (0, 1, 2, …, *n*), *N* stands for total number of observed synonymous codons (*N* ≥ 0), and μ is the genomic mean (0 < μ < 1). The link function for binomially distributed values is *f*(*n*) = *log*[*n*/(1-*n*)].

When the maximum likelihood estimation was converged, codon counts were transformed to Z values using the Wald-test (**Fig. S1**; summary.gamlss function of the gamlss package). In specific cases of *n* = 0 and *n* = *N*, the Z value was set to the maximum value among the other ORFs (**Table S1B**). The pBI function of gamlss package was then used to estimate P values, after which a type I error correction (i.e., false positive error) was performed using the qvalue package^37^ to calculate q-values (analog of false discovery rate, FDR; **Table S1C**). ORFs with at least one codon significantly deviating from the genome average (q < 0.05) were considered as codon-biased ORFs.

### Codon clustering and functional enrichment

Dimensionality of the obtained codon bias profile was reduced by principal component analysis (PCA; prcomp function of stats package). Then, first two PC spaces (cumulative contribution ratio (CCR) = 38.09%; **Fig. S2B,C**), were subject to Gaussian mixture model clustering using Mclust function of mclust package.^38^

Cluster-specific codon-biased ORFs in each cluster (**Fig. 2B**), including 371, 133, 286, and 284 ORFs for cluster I to IV, respectively, were used for GO enrichment analysis. Fisher’s exact test (one-sided, fisher.test function of R stat package) at FDR = 0.05 (qvalue package) was employed to assign GO terms to each cluster.

### Generating the functional profile

To generate the function profile, the basic version of the GO (go-basic.obo) and gene annotations were downloaded from Gene Ontology Consortium (http://geneontology.org/docs/download-ontology/) and SGD, respectively. Then, a Boolean matrix of GO terms was generated in which if an ORF was annotated by a GO, the value was TRUE; otherwise, FALSE. Next, a GO slimmer process was done in five steps: 1) removing global GO terms (*i.e.*, >200 repetitions), 2) removing GO terms with identical sets of annotated ORFs, 3) removing non-codon-biased ORFs, 4) removing unique GO terms (i.e., <2 repetitions), and 5) removing ORFs with no annotations. The obtained matrix served as the functional profile (including *m* codon-biased ORFs × *n* GO terms) for further analysis.

#### Clustering of the function profile

To cluster codon-biased ORFs into functional clusters with no common term, first binary distances were measured (dist function of stats package) between each pair, using the functional profile. Second, hierarchical clustering analysis was applied to the obtained distance matrix (complete linkage method; hclust function of stats package). Clusters were then defined by “static branch cutting” at a height <0.99, which yielded 137 clusters. Finally, clusters with >2 members (115 of 137 clusters) were considered for further analysis (**Table S5**).

Next, the obtained functional clusters were used for GO enrichment analysis. Fisher’s exact test (one-sided, fisher.test function of stat package; FDR = 0.05) was employed to assign the most appropriate GO terms to each cluster. If more than one GO term was enriched to a cluster, the one with the lowest P value was selected as the representative function (**Table S5**).

#### Construction of the functional network: Dimensionality reduction

A two-step strategy, including PCA followed by canonical correlation analysis (CCA), was designed to obtain linear combinations of codon bias and GO profiles (**Fig. S4**). First, the profile of codon bias was subjected to PCA (prcomp function of stats package). Hereinafter, PCs accounting for CCR = 95% are referred to as codon bias principal components (cPCs). Then, PCA was applied to the functional profile. According to De Haan *et al*.,^39^ applying PCA to a Boolean matrix of GO annotations reduces the dimensionality while preserving the structure of the functional relationships among the records. Hereinafter, PCs accounting for CCR = 99% are referred to as GO principal components (gPCs).

Next, CCA was applied to cPCs and gPCs (cancor function of stats package). The significance of the canonical variables was tested using Bartlett’s Chi-square test (FDR = 0.05) for both codon bias canonical variables (cCV) and GO canonical variables (gCV).

#### Construction of the functional network: Capturing combinatorial optimization

To detect influential factor(s) for differentiating functional clusters, multiple logistic regression analysis was applied between each pair of functional clusters (*_p_*C_2_; *p* is number of clusters) with combinational optimization techniques for CVs as explanatory variables. Logistic regressions were performed using brglm function.^40^ After adaptation of the brglm function, combinatorial optimization was implemented using the step function (stats package) and the best model was selected based on Akaike’s information criterion (AIC). The selected model was then compared to the null model by likelihood ratio test using lrtest function.^41^

#### Construction of the functional network: Associations among functional clusters

To detect relationships between the functional clusters (with >2 members), a pairwise CCA between every pair (*_p_*C_2_; *p* is number of clusters) was performed using cCVs. First, to avoid over-fitting, PCA was applied to cCV scores of members of each cluster. Then, CCA was applied to the obtained PC scores of every pair (where the cumulative number of PC scores of every pair is less than the number of significant CVs). Finally, the obtained heterogeneous canonical variables (hCVs) served as independent components correlating functional clusters. The significance of the canonical correlation coefficient of the first hCVs was tested using Bartlett’s chi-squared test at FDR = 0.1% (qvalue package).

#### Construction of the functional network: Visualization of codon bias analogy

Pairwise similarities between codon-biased ORFs were calculated as Pearson’s correlation coefficients (*r*) using significant cCVs (FDR = 0.05). Then, codon-biased ORFs were visualized as a network of the obtained *r* values (FDR = 5%) and significant inter-clusters (Bartlett’s chi-squared test; FDR = 0.1%) relations using Spring layout of qgraph function of qgraph package.^42^

#### Analysis of the codon bias and gene-function network

Core clusters were defined based on number of significant inter-clusters relations. Clusters were divided into two groups by applying Poisson mixture-model-based clustering using gamlssMX function of gamlss.mx package.^43^ Finally, clusters with ≥11 associations were considered as core clusters (**Fig. S8A**).

Dense clustering was detected according to the median values of Euclidean distances between the center of a cluster and its member on the two-dimensional network (**Fig. 4**). Clusters were divided into two groups by applying gamma mixture-model-based clustering using gamlssMX function of gamlss.mx package. Finally, clusters with <0.1 of median distance were considered as dense clusters (**Fig. S8B**).

#### Evaluating functional information gain

In order to evaluate functional information gain, this study was compared to our past study^21^ and relative synonymous codon usage (RSCU) analysis, a codon specific metric.^22^ RSCU values were calculated for 59 codons and ORFs with at least one RSCU value greater than mean of ribosomal genes (= 3.41) were considered codon-biased. Precision-recall was calculated as previously described^23^ with minor modifications. The *r* values (cor function of stats package) of all pairs of codon-biased ORFs were calculated using 59 Z values or RSCU values. The pairs were sorted in descending order of *r* values and were ranked accordingly. If each pair was co-annotated by at least one GO term (given the functional profile), it counted as true positives (TPs) for each n^th^ rank of ORF pairs (i.e., recall). Then, each TP was divided by its rank to estimate precision.

## Declarations

### Ethics approval and consent to participate

Not applicable.

### Consent for publication

Not applicable.

### Availability of data and materials

All data used here are presented in **Supplementary Information**. Source codes are available for download at GitHub at https://github.com/dedonlab/ASCS

### Authors’ contributions

F.G., P.C.D., and T.B. conceived the study. F.G. developed data-processing workflows and computational analyses. F.G. implemented the methodology. S.O. and S.B. contributed insights for the method development. F.G., T.B., and P.C.D. drafted the manuscript and all authors contributed to reviewing and revising the manuscript.

## Acknowledgements

The authors thank members of the Dedon and Begley labs for helpful discussions.

## Supplementary Figure Legends

**Figure S1**. Workflow for ASCS functional analysis of codon usage patterns.

**Figure S2. Codon analytics of budding yeast (*S. cerevisiae*; S288C).** (**A**) Bar plot of number of ORFs possessing significant biases in 59 codons (FDR = 0.05). CGC, CGG, and CGT (Arg), CAC (His), and TCC (Ser) are never under-used in all genes. GGT (Gly), TTG (Leu), and AAG (Lys) are the top three over-used codons. (**B**) Bar plot of ratio of 64 codons in the whole genome (5,797 ORFs). 18 codons form ∼50% of the whole genome. These codons are mainly A/T ending codon (inset bar plot on left) while the rest are mostly C/G ending (inset bar plot on right). (**C-E**) Scatter plots of genomic ratio of 59 codons and number of ORFs with under-used codon (**C**), over-used codon (**D**), and both (**E**). Each circle is a codon.

**Figure S3. Multivariate statistical analysis of synonymous codon usage**. (**A**) Principal component analysis (PCA) of Z values of codon bias ORFs. Black bars (left axis) indicate proportion of variance, red circles (right axis) indicate contribution ratio (CCR), and horizontal dashed lines (right axis) indicate CCRs of 60% and 90%. The first two PC scores (CCR = 38.09%) were used for Gaussian mixture model (GMM) clustering. (**B**) Defining the number of components for GMM clustering. The number of underlying Gaussian distributions was defined based on Bayesian information criterion (BIC) values of models with differing parametrizations. A VII-model (spherical distributions with variable volume, equal shapes, and no orientations), see Scrucca et al. (2016), with four components provided the best fit. (**C**) Posterior membership probability heatmap. Each value shows the posterior probability based on the VII-model in “B”. The heatmap was generated using logarithmic transformation of conditional probabilities from expectation maximization. The dendrogram illustrates model-based hierarchical agglomerative clustering based on the Gaussian probability model for maximizing the resulting likelihood. (**D**) Logos of members of each cluster.

**Figure S4. Clustering of the functional profiles of codon-biased ORFs.** Dendrogram of hierarchical clustering analysis (binary distances and complete linkage method) of functional profiles of codon-biased ORFs. Clusters are defined by “static branch cutting” at a height <0.99. Examples of 18 clusters are displayed with corresponding cluster numbers (Table S5). Numbers in parentheses represent the number of members in the cluster.

**Figure S5. An eye diagram of the canonical correlation analysis (CCA) procedure for drawing linear combinations between functional and codon-biased profiles.** Separate principal component analyses (PCA) of the ORFs for GO terms (gPCs) and codon biases (cPCs) followed by CCA resulted in the extraction of 11 linear pairs of canonical variables (CVs) connecting the codon-bias profile (cCVs) to the GO functional profile (gCVs). Edges show significant loadings (*P* < 0.05, Student t-test) with more than one relationship to other nodes. gPCs: GO terms Principal Components; gCVs: GO terms Canonical Variables; cCVs: codon- bias Canonical Variables; cPCs: codon-bias Principal Components.

**Figure S6. Capturing the best combination of codon-bias canonical variable (cCV) space to differentiate functional clusters**. (**A**) Heatmap of number of required CV space(s) to separate functional clusters in a pairwise manner (*_p_*C_2_; *p* is number of clusters) using logistic regressions followed by a stepwise model selection based on AIC. Not-significant means the null model (a model that includes only an intercept) was chosen as the final model. **Inset**: Bar plot of frequency of required CVs. (**B-C**) Scatter plots of biased distributions of functional clusters in codon-biased canonical variable (cCV) spaces. Members of cluster 20 and cluster 42 can be best separated using cCV1, cCV3, and cCV10 (**B**). cCV1 and cCV7 are required to distinguish members of cluster 14 from cluster 42 (**C**). cCV: codon-bias Canonical Variable.

**Figure S7. Pearson’s correlation analysis between codon-bias canonical variables (cCVs) and Z values of codons.** Bar plots of Pearson’s correlation analysis (*r*) of 11 significant cCVs (Fig. S5) and Z values of 59 codons of codon-bias ORFs. The highest *r* value in each space is shown in gray.

**Figure S8. Unraveling characteristics of functional clusters.** (**A**) Histogram of the number significant correlation(s) among functional groups (Bartlett’s chi-squared test at FDR = 0.1%). Green and black curves define the underlying populations according to the Poisson mixture-model-based clustering. “Core” groups are defined as having ≥ 11 associations (dashed blue line). (**B**) Histogram of median of Euclidean distances between the center of a cluster and its member on the two-dimensional network in Figure 4. Green and black curves define 2 underlying populations according to the gamma mixture-model-based clustering. “Dense” groups are defined as <0.1 of the median distance from the center of the cluster (dashed blue line). (**C**) A graphical representation of codon-bias ORFs and their functions is shown (Spring layout; Figure 4). Of the 115 functional groups (Table S5), 10 (orange and turquoise dots) and 63 (red and turquoise dots) were identified as core and dense groups, respectively. Additionally, four groups (turquoise dots) were identified as both core and dense groups. Other clusters (black dots) did not fall into any of these categories.

**Figure S9. Associations among functional groups.** (**A**) The bar plot displays the number of associations for each functional group, with “core” clusters (≥11 associations; Fig. S9A) are shown in orange. GO terms assigned to the core clusters are shown as “Cluster number. GO term description (number of members)”. See Table S5 for the GO categories. The heatmap shows significant canonical correlation coefficients (red squares) of the first heterogeneous canonical variables (hCV1) of pairwise CCA (Bartlett’s chi-squared test at FDR = 0.1%). (**B**) The dendrogram represents the hierarchical cluster analysis of all hCV1s (“1 – *r*” and average linkage).

**Figure S10. Benchmarking of the biological information gain.** (**A**) Venn diagram of codon-bias ORFs obtained by this study versus our previous study^1^ and relative synonymous codon usage (RSCU; a codon specific metric). (**B**) Precision-recall plot for positive correlation coefficients^2^. TP: True Positive; FP: False Positives. (**C-D**) Global network of functional relationships among codon-bias ORFs of this study (**C**) and Begley et al. (2007; **D**).

**Figure S11. Mapping H_2_O_2_ response genes onto the codon-bias functional network.** (**A**) A graphical representation of codon-bias ORFs and their functions is shown (Spring layout; Figure 4). Genes for up- and down-regulated proteins under H_2_O_2_ stress^3^ are shown in red and green, respectively in each functional cluster. Other members in each cluster are shown in black. (**B**) Bar plots of number of genes for up- and down-regulated proteins in each functional cluster.

**Figure S12. A comparison of gene sequence similarity and similarity of codon usage patterns in genes encoding large and small ribosome proteins in *S. cerevisiae,* including gene paralogs.** Insets: Upper plot is a blown-up of a section of the main graph, showing gene comparisons with *r* ≥ 0.5 including the paralogous gene pairs. The red line represents a simple regression line. The lower plot shows the residuals of the regression line in the upper plot.

## Footnotes

1 Effect size is the median value of the set of absolute Z values of members in each cluster, with larger Z values reflecting stronger codon biases. The set of metabolic clusters with effect size >80: Cluster 73, hexose metabolic process, 80.9; Cluster 8, sulfur compound metabolic process, 82.9; Cluster 29, alcohol metabolic process, 86.9; Cluster 2, nucleotide metabolic process, 88.9

