## Supplementary Information for "Synonymous codon usage defines functional gene families"

**Contents**

**Figure S1**. Workflow for ASCS functional analysis of codon usage patterns.

**Figure S2**. Codon analytics of budding yeast (*S. cerevisiae*; S288C).

**Figure S3**. Multivariate statistical analysis of synonymous codon usage.

**Figure S4.** Clustering of the functional profiles of codon-biased ORFs.

**Figure S5.** An eye diagram of the canonical correlation analysis (CCA) procedure for drawing linear combinations between functional and codon-biased profiles.

**Figure S6.** Capturing the best combination of codon-bias canonical variable (cCV) space to differentiate functional clusters.

**Figure S7.** Pearson’s correlation analysis between codon-bias canonical variables (cCVs) and Z values of codons.

**Figure S8.** Unraveling characteristics of functional clusters.

**Figure S9.** Associations among functional groups.

**Figure S10.** Benchmarking of the biological information gain.

**Figure S11.** Mapping H_2_O_2_ response genes on to the codon-bias functional network.

**Figure S12.** A comparison of gene sequence similarity and similarity of codon usage patterns in genes encoding large and small ribosome proteins in *S. cerevisiae*.

**Tables S1-S7**. Provided as a separate Excel spreadsheet

**Supplementary References**

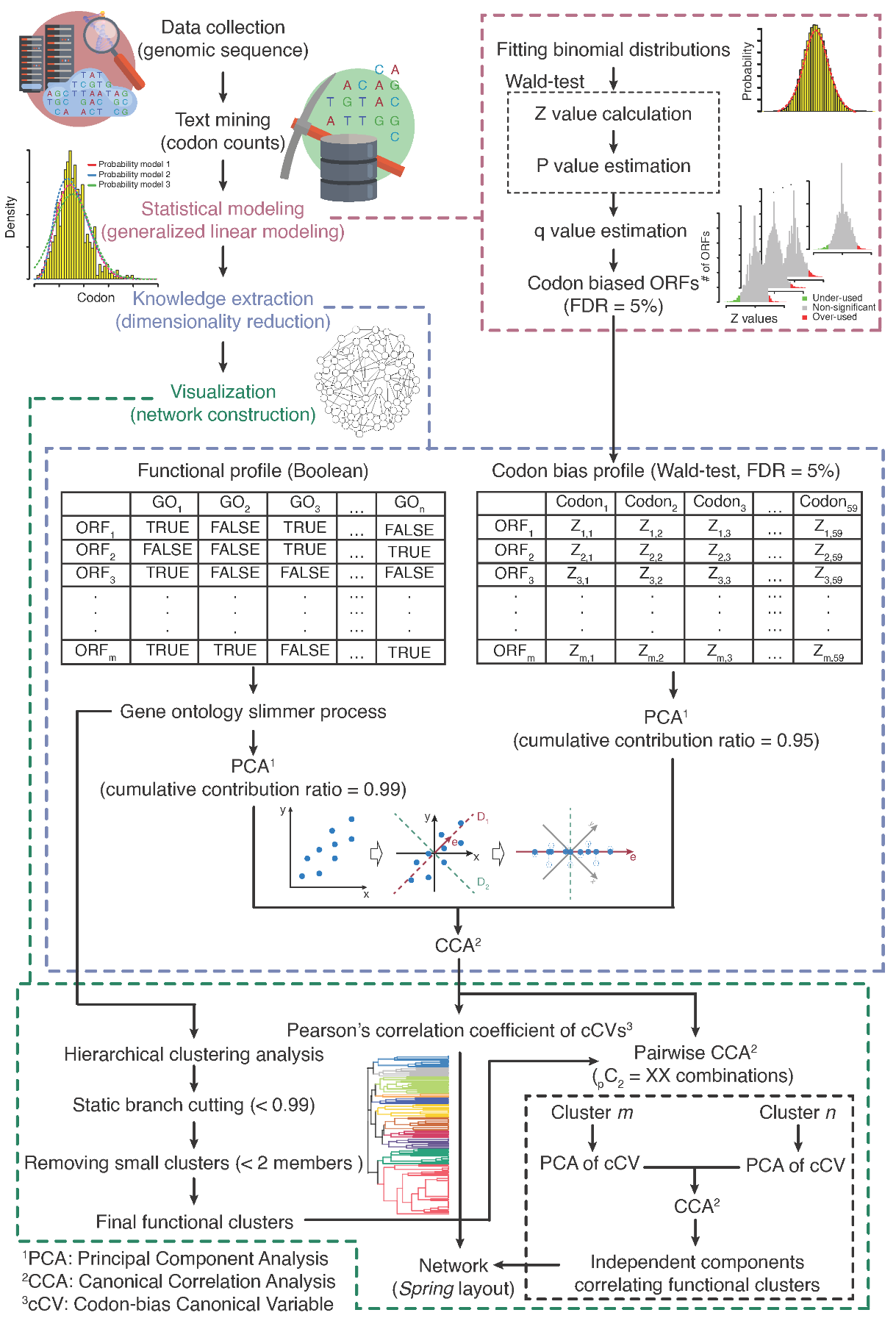

**Figure S1**. Workflow for ASCS functional analysis of codon usage patterns.

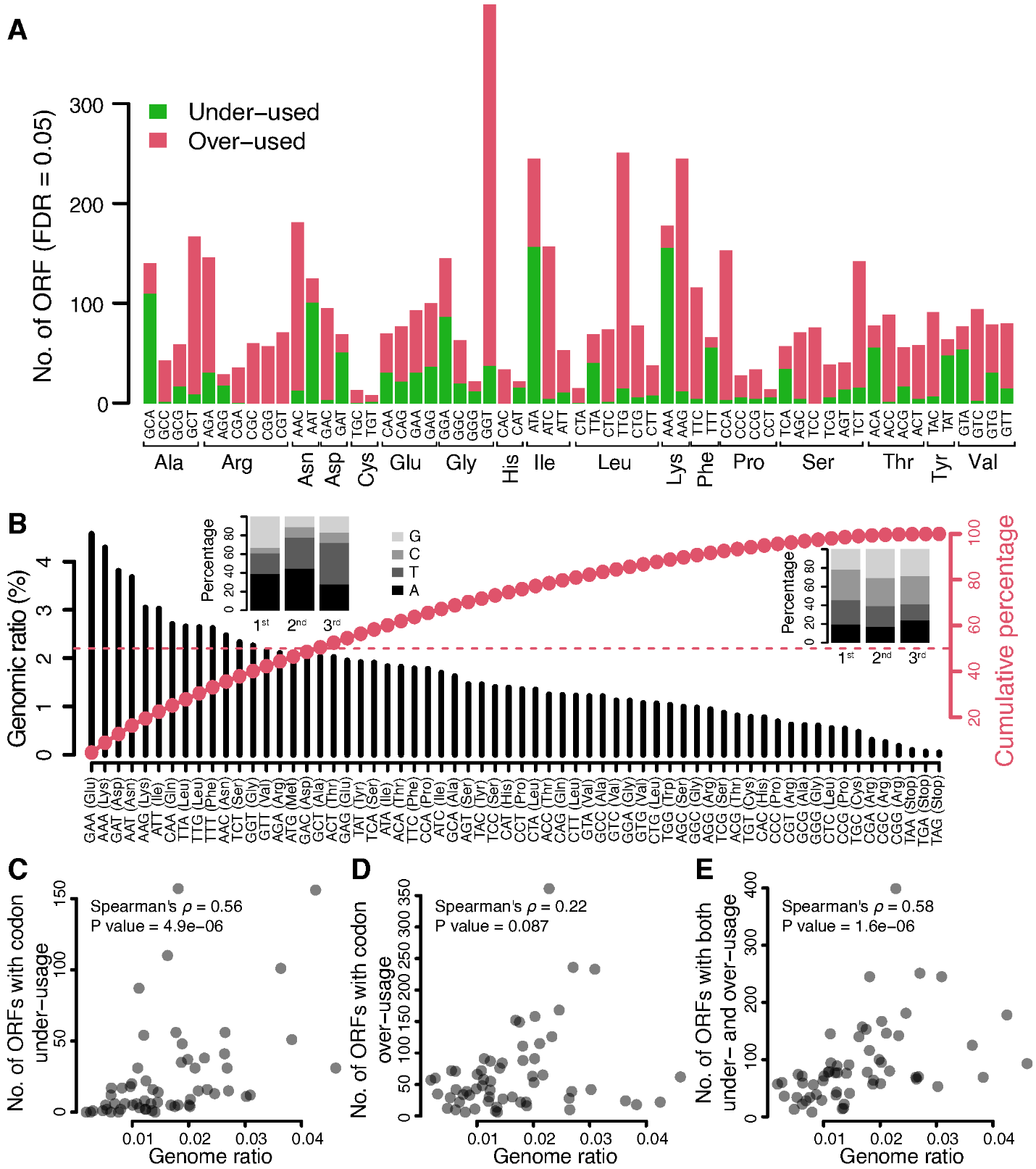

**Figure S2. Codon analytics of budding yeast (*S. cerevisiae*; S288C).** (**A**) Bar plot of number of ORFs possessing significant biases in 59 codons (FDR = 0.05). CGC, CGG, and CGT (Arg), CAC (His), and TCC (Ser) are never under-used in all genes. GGT (Gly), TTG (Leu), and AAG (Lys) are the top three over-used codons. (**B**) Bar plot of ratio of 64 codons in the whole genome (5,797 ORFs). 18 codons form ~50% of the whole genome. These codons are mainly A/T ending codon (inset bar plot on left) while the rest are mostly C/G ending (inset bar plot on right). (**C-E**) Scatter plots of genomic ratio of 59 codons and number of ORFs with under-used codon (**C**), over-used codon (**D**), and both (**E**). Each circle is a codon.

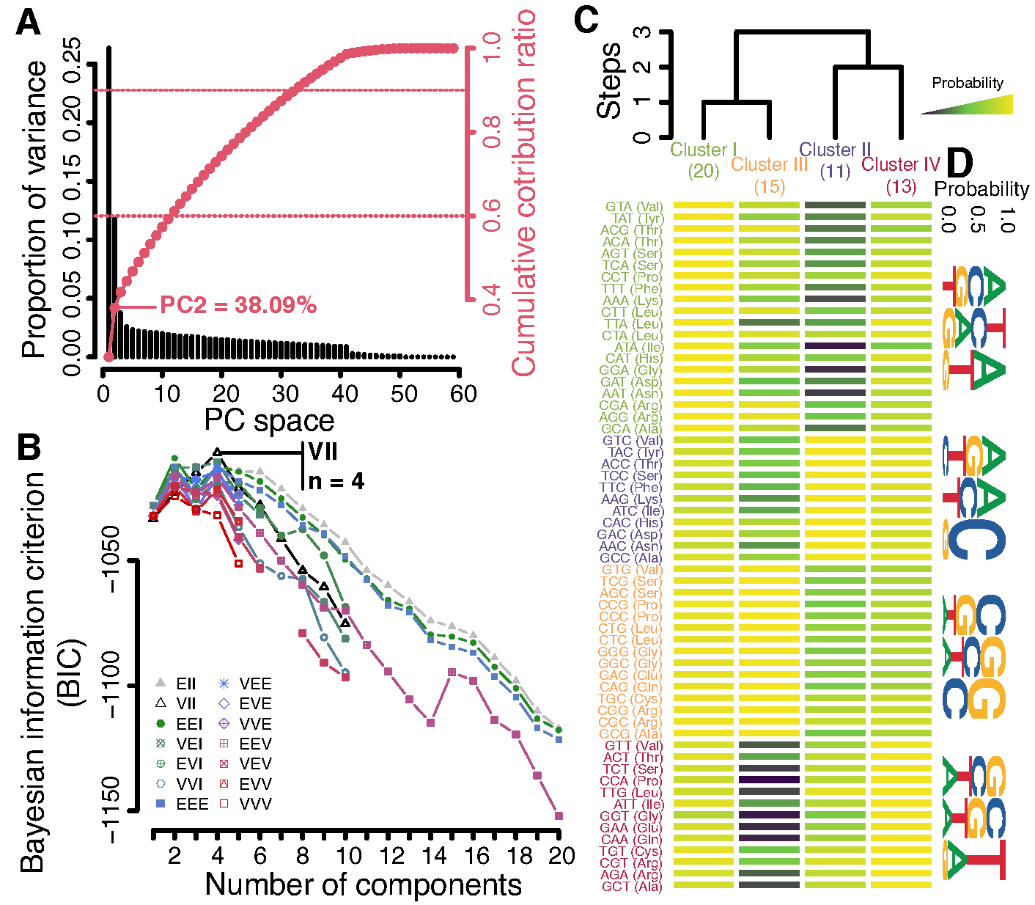

**Figure S3. Multivariate statistical analysis of synonymous codon usage**. (**A**) Principal component analysis (PCA) of Z values of codon bias ORFs. Black bars (left axis) indicate proportion of variance, red circles (right axis) indicate contribution ratio (CCR), and horizontal dashed lines (right axis) indicate CCRs of 60% and 90%. The first two PC scores (CCR = 38.09%) were used for Gaussian mixture model (GMM) clustering. (**B**) Defining the number of components for GMM clustering. The number of underlying Gaussian distributions was defined based on Bayesian information criterion (BIC) values of models with differing parametrizations. A VII-model (spherical distributions with variable volume, equal shapes, and no orientations), see Scrucca et al. (2016), with four components provided the best fit. (**C**) Posterior membership probability heatmap. Each value shows the posterior probability based on the VII-model in “B”. The heatmap was generated using logarithmic transformation of conditional probabilities from expectation maximization. The dendrogram illustrates model-based hierarchical agglomerative clustering based on the Gaussian probability model for maximizing the resulting likelihood. (**D**) Logos of members of each cluster.

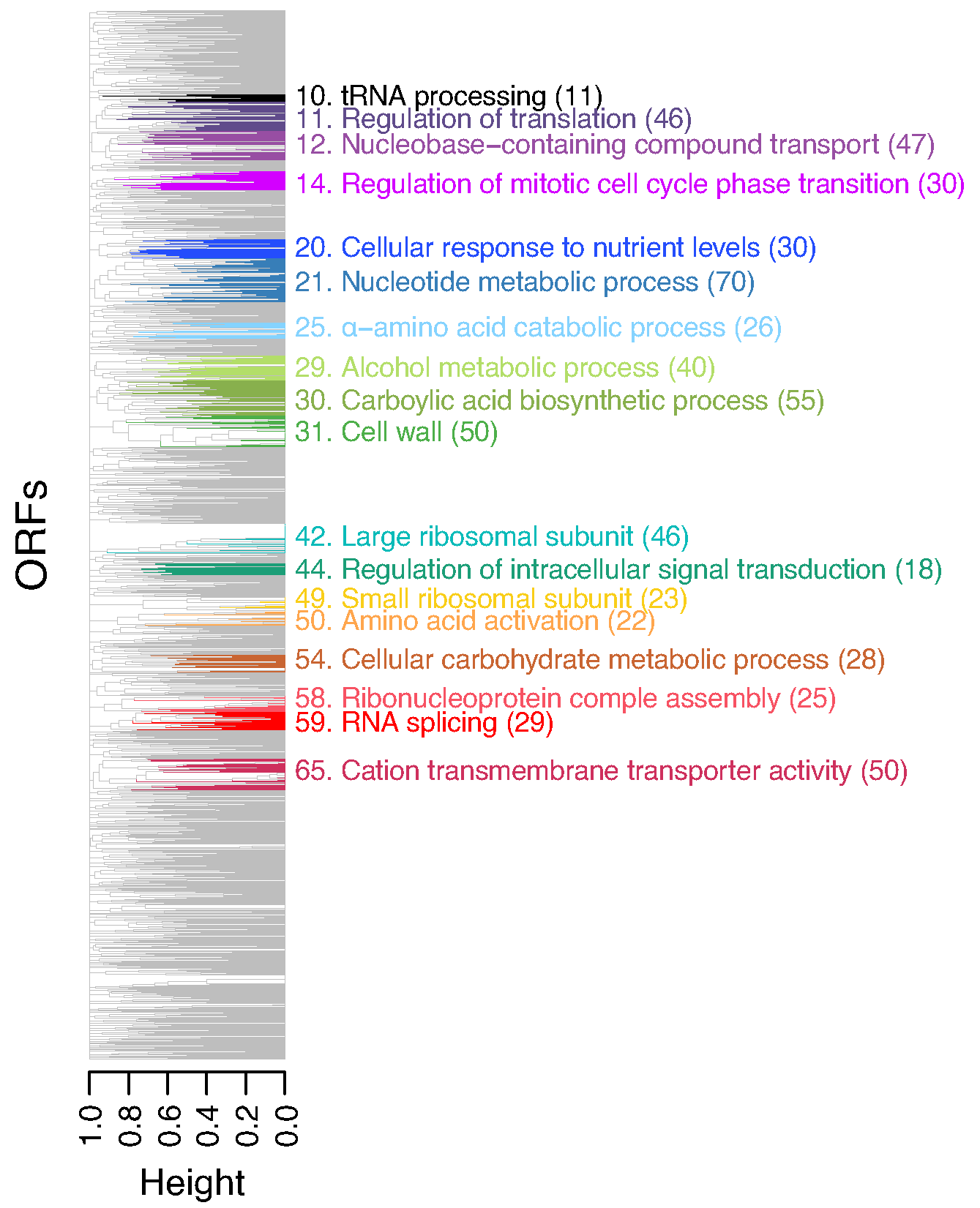

**Figure S4. Clustering of the functional profiles of codon-biased ORFs.** Dendrogram of hierarchical clustering analysis (binary distances and complete linkage method) of functional profiles of codon-biased ORFs. Clusters are defined by “static branch cutting” at a height <0.99. Examples of 18 clusters are displayed with corresponding cluster numbers (Table S5). Numbers in parentheses represent the number of members in the cluster.

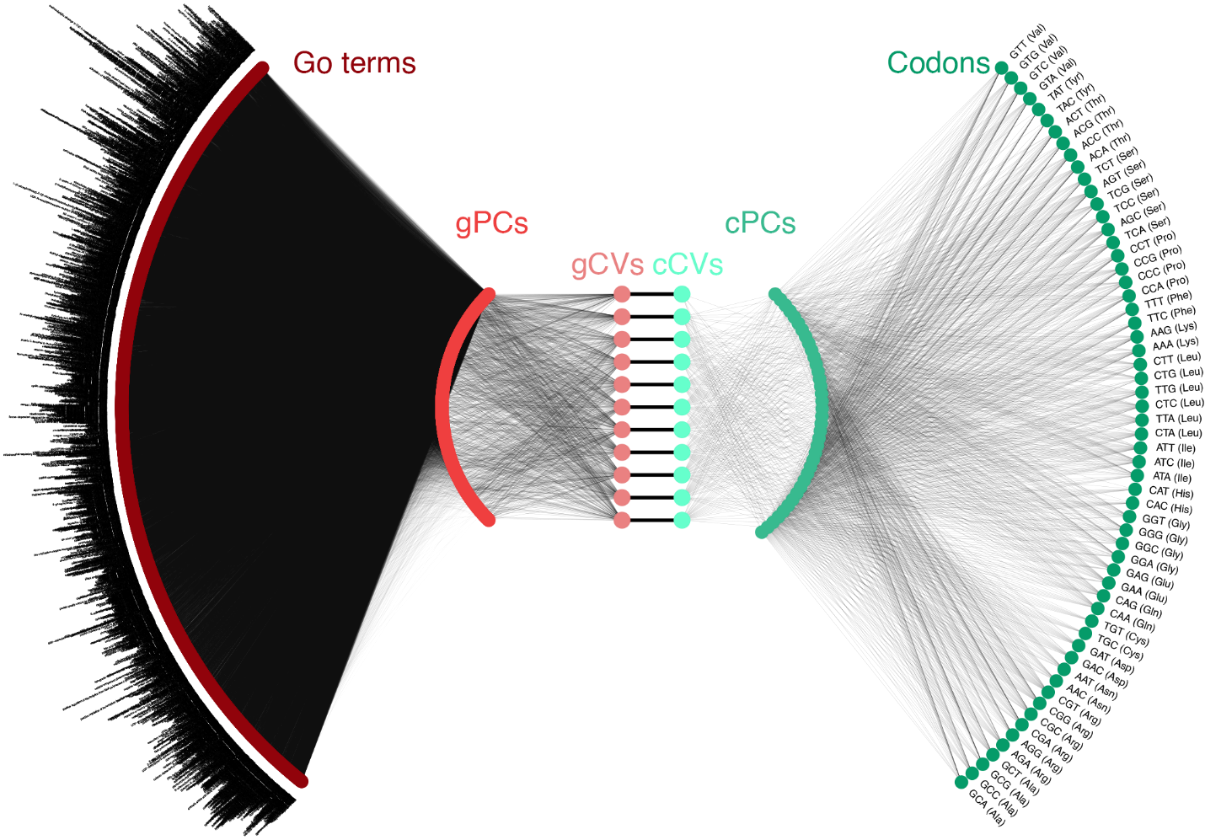

**Figure S5. An eye diagram of the canonical correlation analysis (CCA) procedure for drawing linear combinations between functional and codon-biased profiles.** Separate principal component analyses (PCA) of the ORFs for GO terms (gPCs) and codon biases (cPCs) followed by CCA resulted in the extraction of 11 linear pairs of canonical variables (CVs) connecting the codon-bias profile (cCVs) to the GO functional profile (gCVs). Edges show significant loadings (*P* < 0.05, Student t-test) with more than one relationship to other nodes. gPCs: GO terms Principal Components; gCVs: GO terms Canonical Variables; cCVs: codon-bias Canonical Variables; cPCs: codon-bias Principal Components.

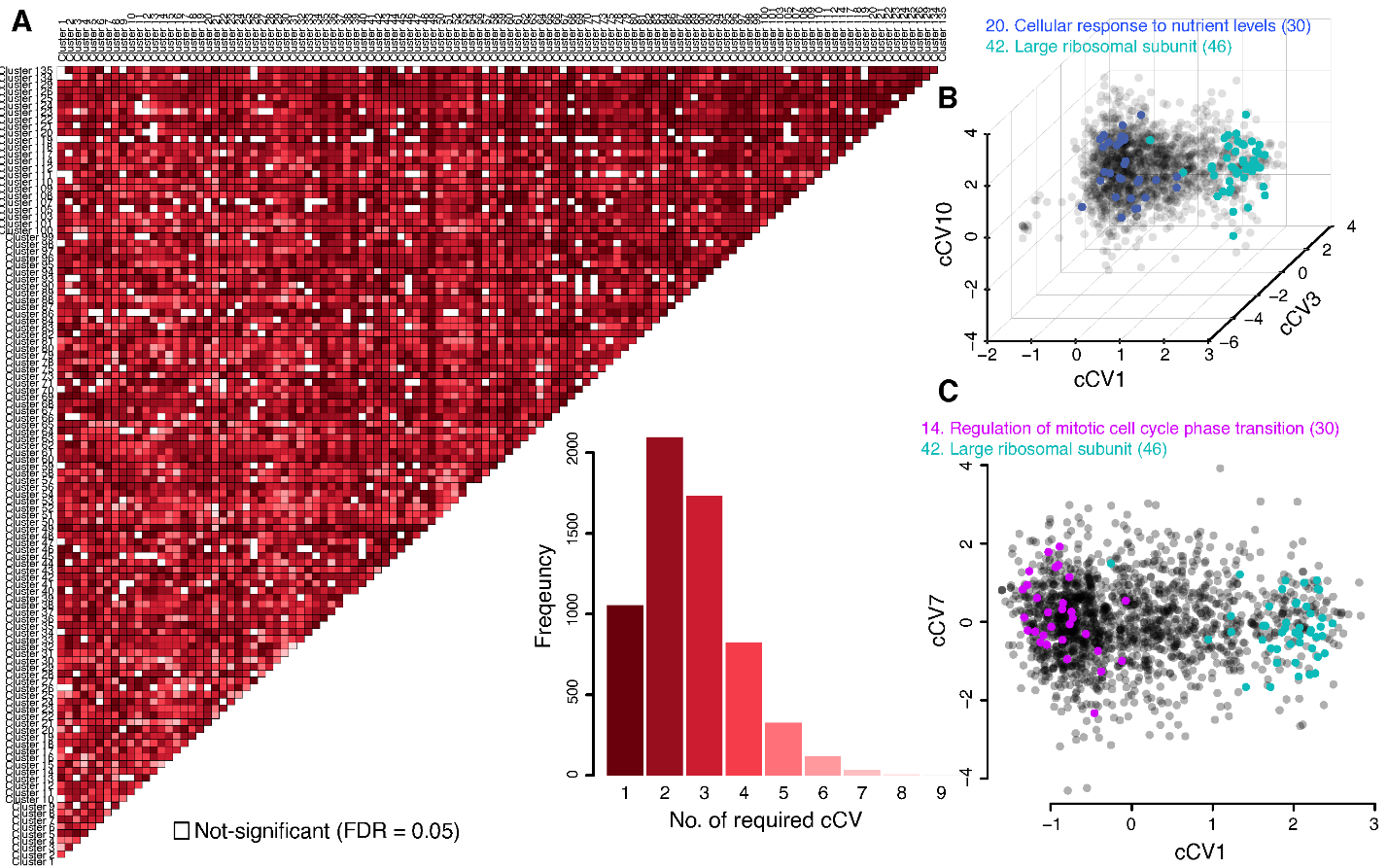

**Figure S6. Capturing the best combination of codon-bias canonical variable (cCV) space to differentiate functional clusters**. (**A**) Heatmap of number of required CV space(s) to separate functional clusters in a pairwise manner (*_p_*C_2_; *p* is number of clusters) using logistic regressions followed by a stepwise model selection based on AIC. Not-significant means the null model (a model that includes only an intercept) was chosen as the final model. **Inset**: Bar plot of frequency of required CVs. (**B-C**) Scatter plots of biased distributions of functional clusters in codon-biased canonical variable (cCV) spaces. Members of cluster 20 and cluster 42 can be best separated using cCV1, cCV3, and cCV10 (**B**). cCV1 and cCV7 are required to distinguish members of cluster 14 from cluster 42 (**C**). cCV: codon-bias Canonical Variable.

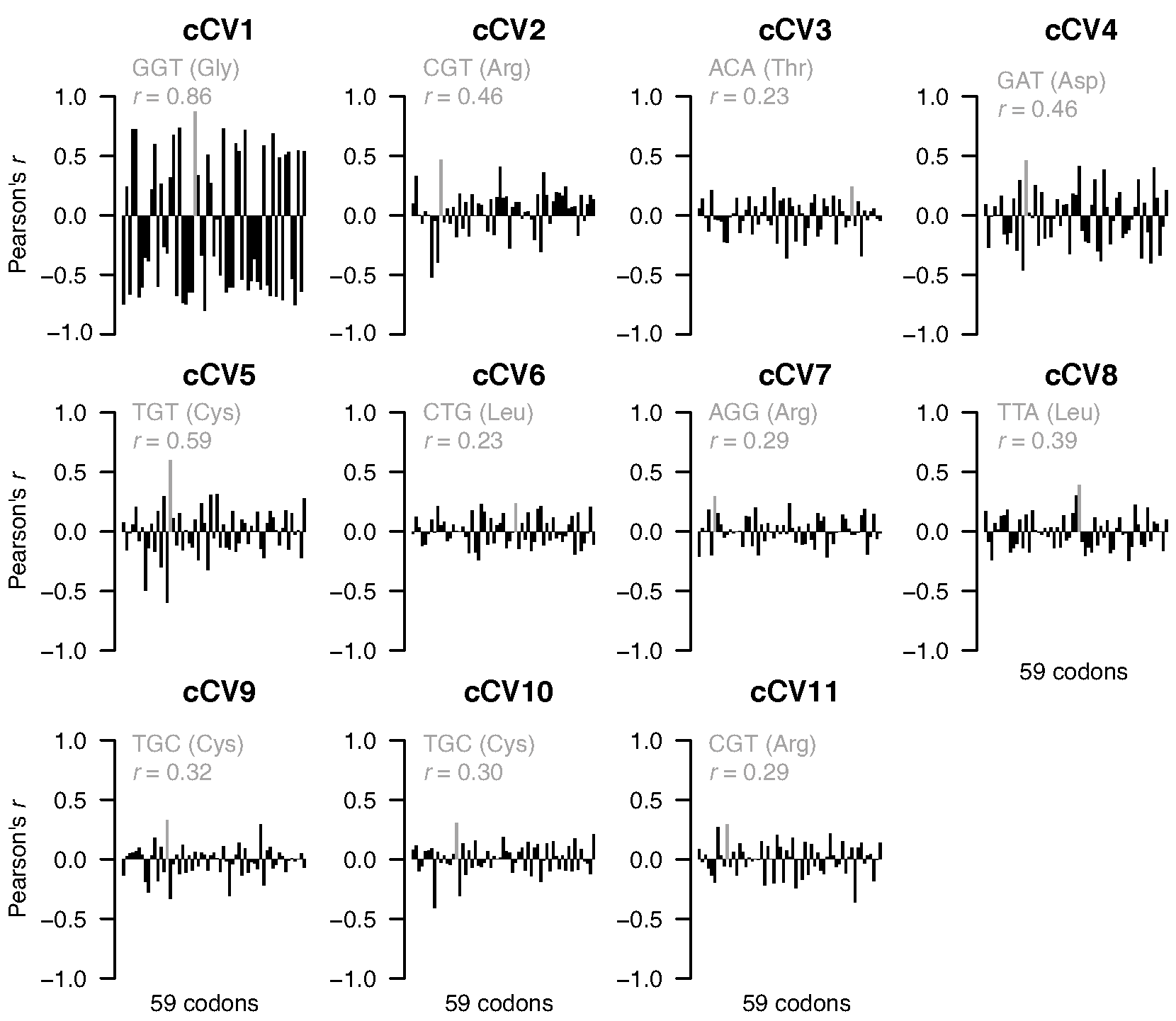

**Figure S7. Pearson’s correlation analysis between codon-bias canonical variables (cCVs) and Z values of codons.** Bar plots of Pearson’s correlation analysis (*r*) of 11 significant cCVs (Fig. S5) and Z values of 59 codons of codon-bias ORFs. The highest *r* value in each space is shown in gray.

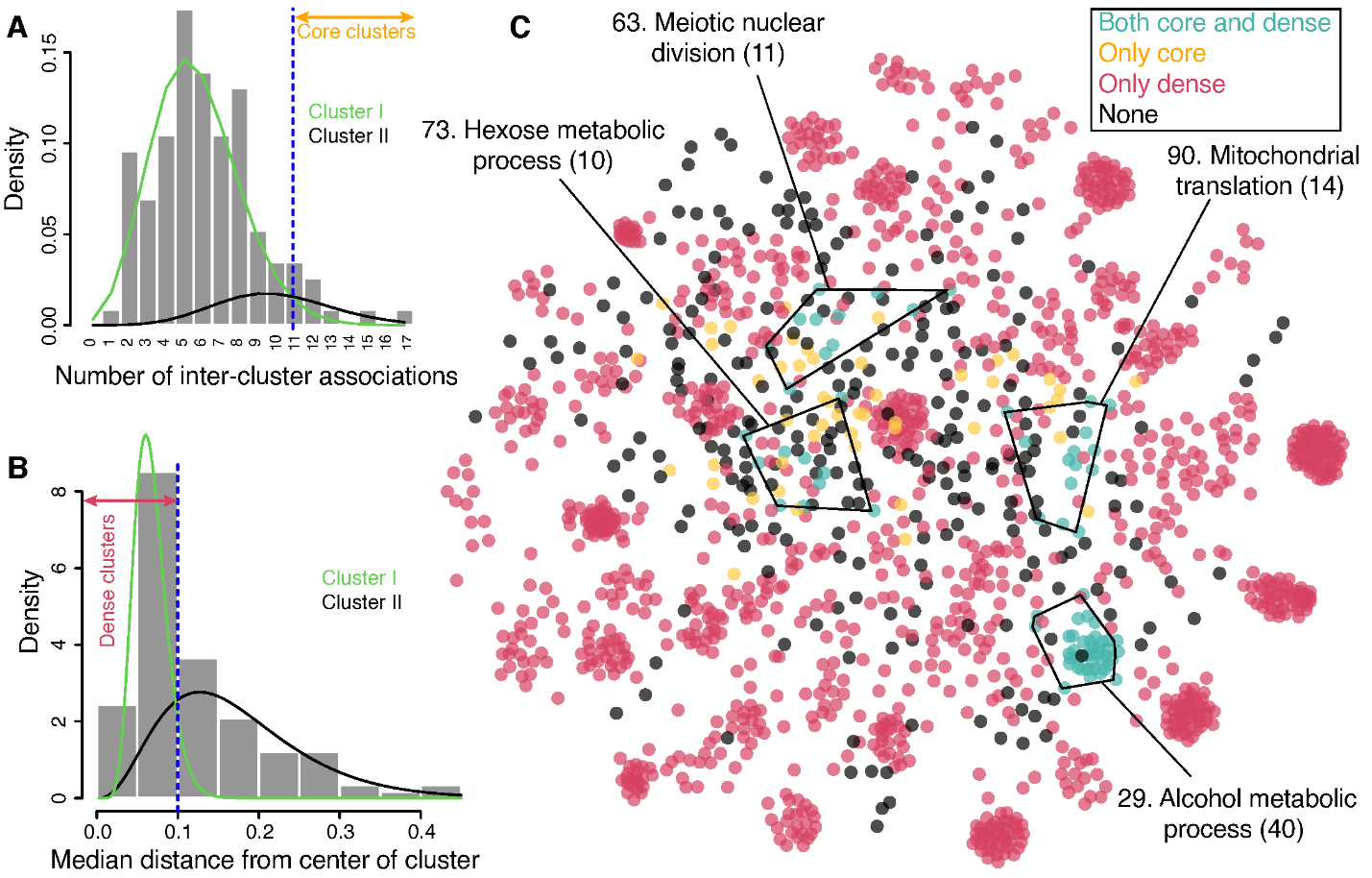

**Figure S8. Unraveling characteristics of functional clusters.** (**A**) Histogram of the number significant correlation(s) among functional groups (Bartlett’s chi-squared test at FDR = 0.1%). Green and black curves define the underlying populations according to the Poisson mixture-model-based clustering. “Core” groups are defined as having ≥ 11 associations (dashed blue line). (**B**) Histogram of median of Euclidean distances between the center of a cluster and its member on the two-dimensional network in Figure 4. Green and black curves define 2 underlying populations according to the gamma mixture-model-based clustering. “Dense” groups are defined as <0.1 of the median distance from the center of the cluster (dashed blue line). (**C**) A graphical representation of codon-bias ORFs and their functions is shown (Spring layout; Figure 4). Of the 115 functional groups (Table S5), 10 (orange and turquoise dots) and 63 (red and turquoise dots) were identified as core and dense groups, respectively. Additionally, four groups (turquoise dots) were identified as both core and dense groups. Other clusters (black dots) did not fall into any of these categories.

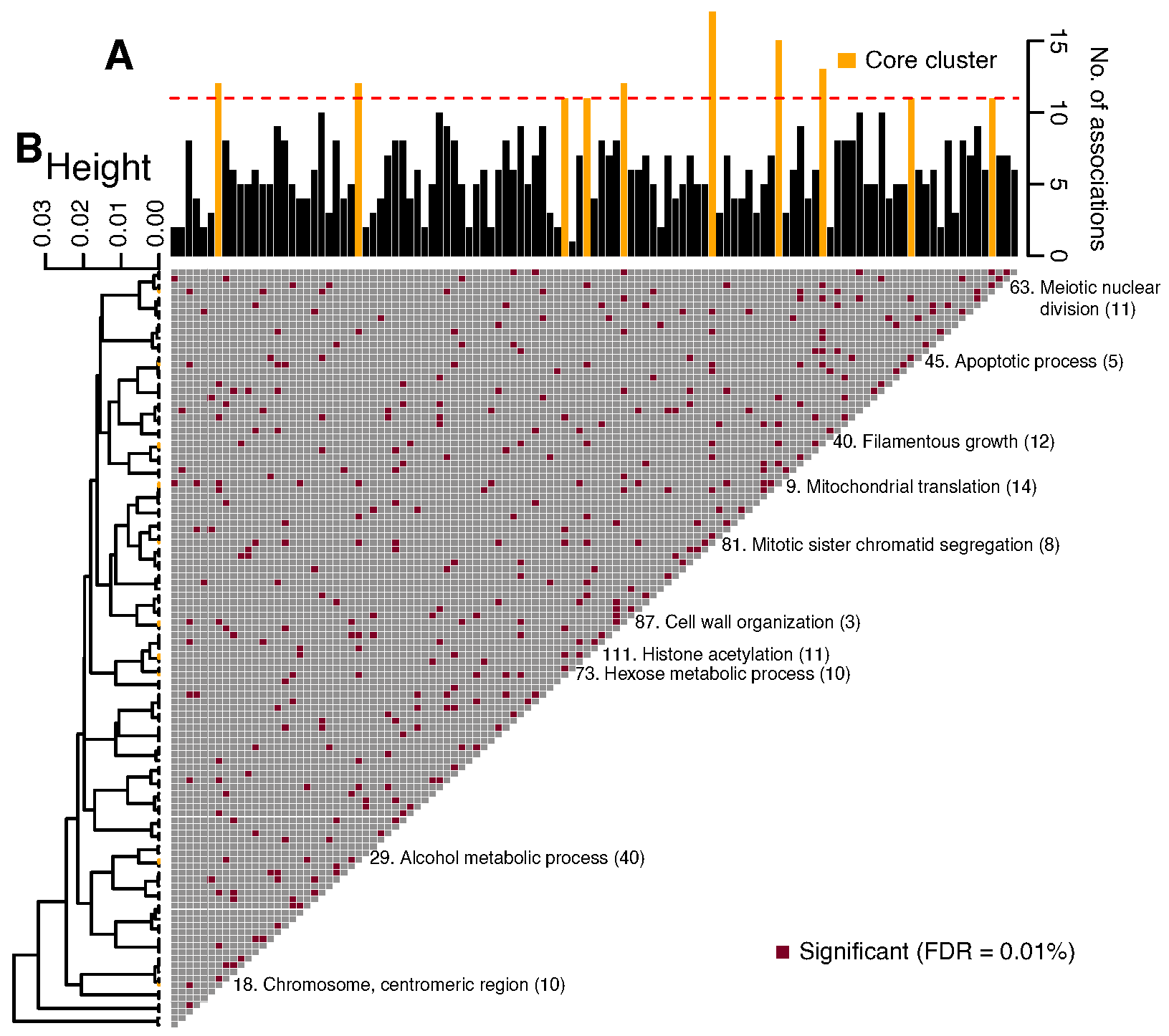

**Figure S9. Associations among functional groups.** (**A**) The bar plot displays the number of associations for each functional group, with “core” clusters (≥11 associations; Fig. S9A) are shown in orange. GO terms assigned to the core clusters are shown as “Cluster number. GO term description (number of members)”. See Table S5 for the GO categories. The heatmap shows significant canonical correlation coefficients (red squares) of the first heterogeneous canonical variables (hCV1) of pairwise CCA (Bartlett’s chi-squared test at FDR = 0.1%). (**B**) The dendrogram represents the hierarchical cluster analysis of all hCV1s (“1 – *r*” and average linkage).

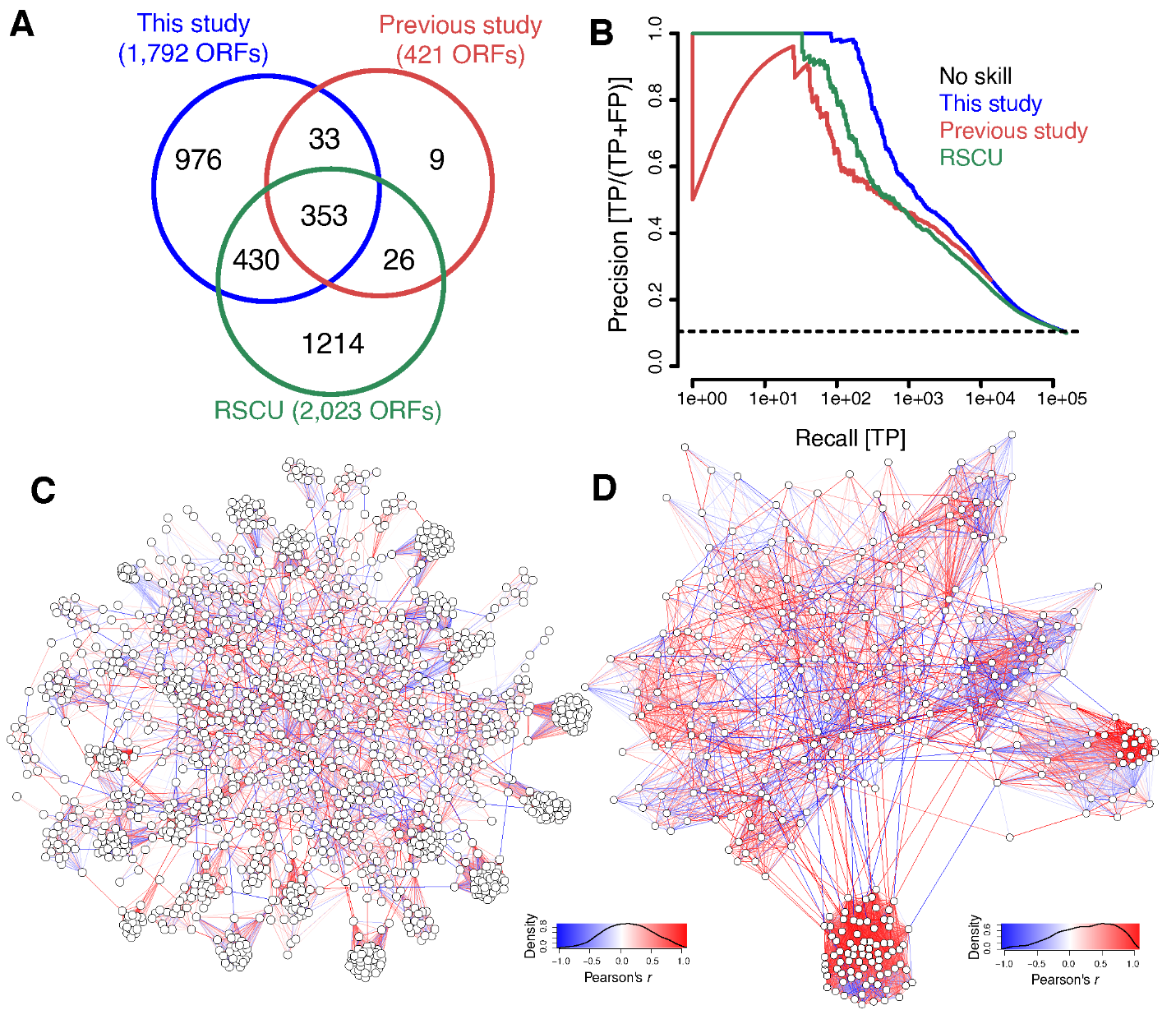

**Figure S10. Benchmarking of the biological information gain.** (**A**) Venn diagram of codon-bias ORFs obtained by this study versus our previous study^1^ and relative synonymous codon usage (RSCU; a codon specific metric). (**B**) Precision-recall plot for positive correlation coefficients^2^. TP: True Positive; FP: False Positives. (**C-D**) Global network of functional relationships among codon-bias ORFs of this study (**C**) and Begley et al. (2007; **D**).

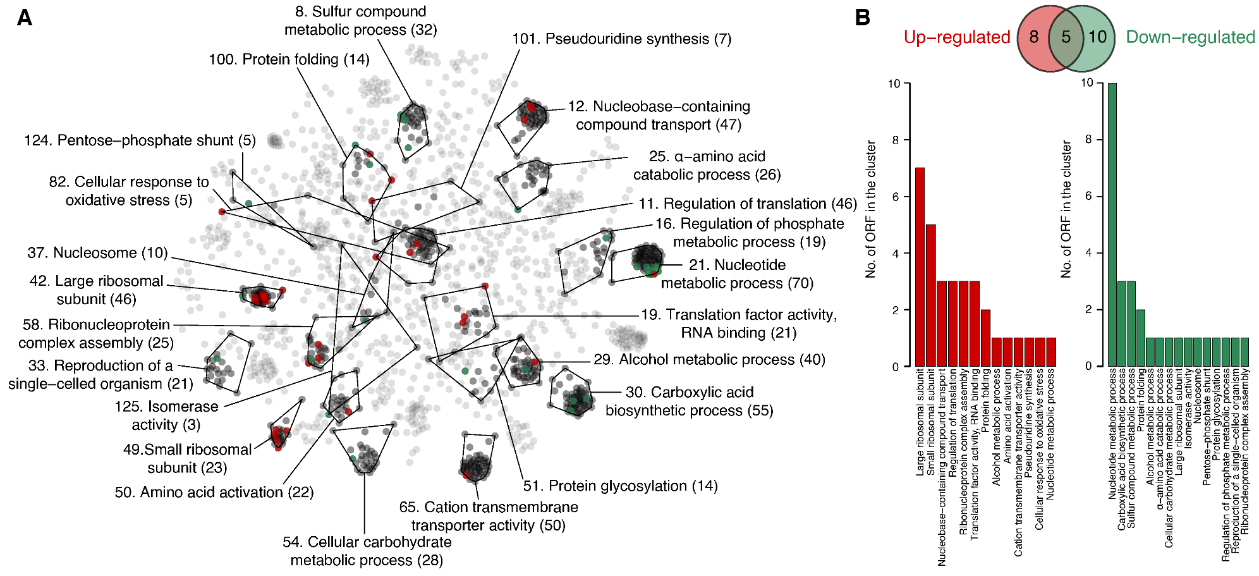

**Figure S11. Mapping H_2_O_2_ response genes onto the codon-bias functional network.** (**A**) A graphical representation of codon-bias ORFs and their functions is shown (Spring layout; Figure 4). Genes for up- and down-regulated proteins under H_2_O_2_ stress^3^ are shown in red and green, respectively in each functional cluster. Other members in each cluster are shown in black. (**B**) Bar plots of number of genes for up- and down-regulated proteins in each functional cluster.

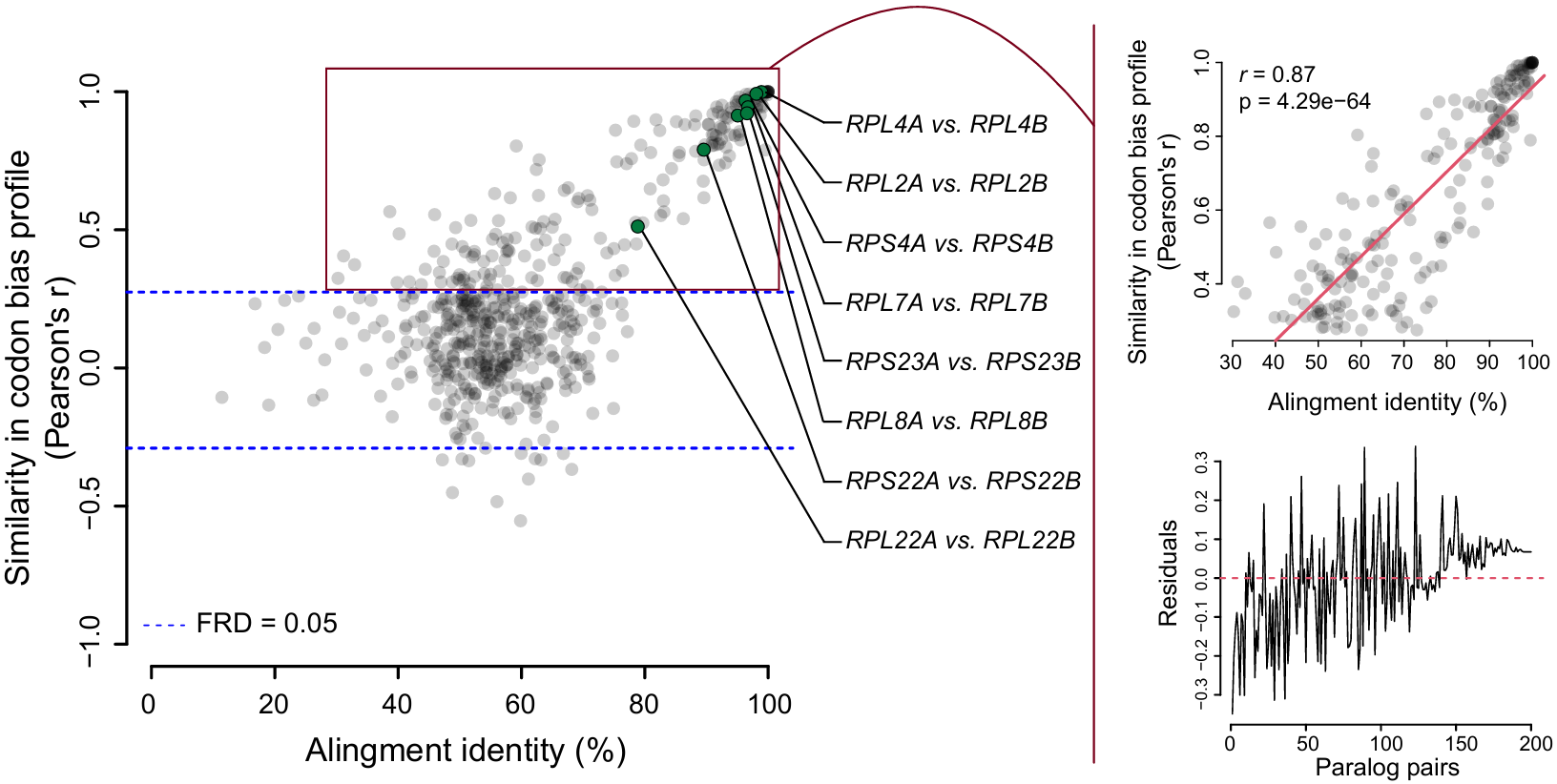

**Figure S12. A comparison of gene sequence similarity and similarity of codon usage patterns in genes encoding large and small ribosome proteins in *S. cerevisiae,* including gene paralogs. Insets:** Upper plot is a blown-up of a section of the main graph, showing gene comparisons with *r* ≥ 0.5 including the paralogous gene pairs. The red line represents a simple regression line. The lower plot shows the residuals of the regression line in the upper plot.
